# A forest model intercomparison framework and application at two temperate forests along the East Coast of the United States

**DOI:** 10.1101/464578

**Authors:** Adam Erickson, Nikolay Strigul

## Abstract

Forest models often reflect the dominant management paradigm of their time. Until the late 1970s, this meant sustaining yields. Following landmark work in forest ecology, physiology, and biogeochemistry, the current generation of models is further intended to inform ecological and climatic forest management in alignment with national biodiversity and climate mitigation targets. This has greatly increased the complexity of models used to inform management, making them difficult to diagnose and understand. State-of-the-art forest models are often complex, analytically intractable, and computationally-expensive, due to the explicit representation of detailed biogeochemical and ecological processes. Different models often produce distinct results while predictions from the same model vary with parameter values. In this project, we developed a rigorous quantitative approach for conducting model intercomparisons and assessing model performance. We have applied our original methodology to compare two forest biogeochemistry models, the Perfect Plasticity Approximation with Simple Biogeochemistry (PPA-SiBGC) and Landscape Disturbance and Succession with Net Ecosystem Carbon and Nitrogen (LANDIS-II NECN). We simulated past-decade conditions at flux tower sites located within Harvard Forest, MA, USA (HF-EMS) and Jones Ecological Research Center, GA, USA (JERC-RD). We mined field data available for both sites to perform model parameterization, validation, and intercomparison. We assessed model performance using the following time-series metrics: net ecosystem exchange, aboveground net primary production, aboveground biomass, C, and N, belowground biomass, C, and N, soil respiration, and, species total biomass and relative abundance. We also assessed static observations of soil organic C and N, and concluded with an assessment of general model usability, performance, and transferability. Despite substantial differences in design, both models achieved good accuracy across the range of pool metrics. While LANDIS-II NECN showed better fidelity to interannual NEE fluxes, PPA-SiBGC indicated better overall performance for both sites across the 11 temporal and 2 static metrics tested (HF-EMS 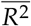 = 0.73, +0.07, 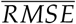 = 4.84, −10.02; JERC-RD 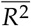 = 0.76, +0.04, 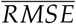 = 2.69, −1.86). To facilitate further testing of forest models at the two sites, we provide pre-processed datasets and original software written in the R language of statistical computing. In addition to model intercomparisons, our approach may be employed to test modifications to forest models and their sensitivity to different parameterizations.

## 1. Introduction

For millenia, timber harvest for economic, militaristic, and social gain was the primary – if not sole – objective of forestry. This focus changed only slightly in the 18^th^ century with the emergence of sustained-yield forest management in Leipzig, Germany (then within the Electorate of Saxony, Holy Roman Empire) [1]. For the first time, controlling the effects of management intensity on land productivity over time was given primary consideration. This followed a history of deforestation extending back to the loss of *Cedrus* forests across the Middle East, as described in the *Epic of Gilgamesh* in the third millennium BCE [2,3]. While sustained-yield forest management was designed to maximize timber production indefinitely, under the spurious assumption that sustained yield is possible solely through *in situ* silvicultural treatments, the concept broadly inspired sustainability science, resilience theory [4], and subsequent work on complex adaptive systems [5].

From its inception, sustainability regarded matters economic, social, and ecological in nature [6]. Yet, economic-focused timber production likely accelerated with increased mechanization in the mid-20^th^ century. As our understanding of abiotic and biotic forest interactions expanded, the core assumptions of stationarity underpinning sustained-yield management lost support. The importance of fire ecology [7], structural complexity [8], trophic interactions [9], and their relation to climate, soil, and ecosystem functioning was soon uncovered. Research on climate impacts on regeneration [10,11] further showed that species compositional changes are likely under current climate trajectories, requiring proactive strategies to sustain yields from extant forests.

Research along this line inspired the concepts of adaptive migration [12], assisted gene flow [13], and precise gene editing of trees with CRISPR/Cas9 [14]. Ecological forestry or sustainable forest management is now the dominant management paradigm, where the focus is on emulating natural processes of succession, disturbance, and migration [15]. Mirroring changes in management, modeling forest ecosystems also underwent a paradigm shift from focusing on sustained yield to ecological forestry and multiple-use management. This has required a remarkable increase in the size and complexity of forest ecosystem models in order to simulate a suite of new complex processes.

Forest models likely began 350 years ago in China with yield tables known as the Lung Ch’uan codes, invented by a women of the Kuo family in Suichuan county, Jiangxi [16]. It was not until the 20^th^ century that the first complex mathematical models of forests emerged. Long restricted to simple models developed with mechanical calculators, digital computers enabled researchers for the first time to explicitly model forest dynamics. Following the development of matrix models [17] and empirical growth-and-yield models such as Prognosis [18,19], a vast array of gap [20], forest landscape [21–25], and terrestrial biosphere models [26–28] have been developed. Models of forest ecosystems vary substantially in application, abstraction, and process-level detail.

Representation of canopy geometry varies from implicit to a single ‘big-leaf’ and detailed three-dimensional crown and root geometry (e.g., modern gap models such as MAESPA [29] and LES [30]). Models of growth range from simple allometric equations (e.g., growth-and-yield models) to light-use efficiency models [31] and first-principles mechanistic models of photosynthesis [32]. Belowground process models similarly vary in structure, from simple stoichiometric relations to carbon and nitrogen cycling with microbial dynamics to a fully mechanistic representation of energetic and biogeochemical processes based on thermodynamics. Current belowground models vary considerably in their process representation and accuracy, with much improvement left to be made [33]. Most belowground models in use globally rely on a variant of the classical Century model [34,35].

Model specialization and generalization ranges from pure research applications in narrowly defined areas (e.g., [29]) to simulating multiple loosely coupled landscape processes to simulating biogeochemical fluxes throughout the world’s forests. A trade-off is thought to exist between realism, precision, and generality [36], with more detailed models requiring higher parameterization costs. Yet, little is known about the net effects of variation in the structure of these models on the precision and accuracy of their predictions across temporal and spatial scales. While such model intercomparisons are common within classes of models such as terrestrial biosphere models, they are seldom applied to gap or forest landscape models. Models operating at different scales are seldom compared within sites. Yet, much can be learned by comparing models that differ in assumptions and structure.

Existing forest model intercomparison projects, or MIPs, in Europe include the stand-level Intersectoral Impact MIP (ISIMIP) [37] and landscape-level Comparison of Forest Landscape Models (CoFoLaMo) [38], the latter conducted under the European Union Cooperation on Science and Technology (COST) Action FP1304 “Towards robust projections of European forests under climate change” (ProFoUnd). Previous efforts in the United States include the Throughfall Displacement Experiment (TDE) Ecosystem Model Intercomparison Project at Walker Branch Watershed in Oak Ridge, Tennessee [39]. Presently, no other forest model intercomparison project is evident for North America. There is a critical need to conduct ongoing forest biogeochemistry model comparisons in this and other regions of the world in order to establish the regional foundation for robust global C cycle projections. In this work, we aim to begin this process for North America with a comparison of the Perfect Plasticity Approximation with Simple Biogeochemistry (PPA-SiBGC) and Landscape Disturbance and Succession with Net Ecosystem Carbon and Nitrogen (LANDIS-II NECN) models, which provide contrasting model structures for representing stand dynamics.

Modern forest landscape models are the result of five key model development phases, listed in chronological order: (1) growth-and-yield models; (2) fire models; (3) gap models; (4) physiological models; (5) hybrid models combining design principles from each [20,40,41]. Terrestrial biosphere models similarly trace their roots back to early one-dimensional physiological models, with land surface models currently in their third generation and dynamic global vegetation models in their second generation [42]. This latest generation of models was intended to address the lack of explicit representation of vegetation dynamics - a critical source of model uncertainty in future climate scenarios [43]. This inspired the aforementioned forest ecosystem model intercomparisons as well as new terrestrial biosphere model designs based on gap models, bypassing the trade-offs of medium-resolution forest landscape models.

Collectively, these efforts yielded a number of new terrestrial biosphere models based on the classical gap model, including the Lund-Potsdam-Jena General Ecosystem Simulator (LPJ-GUESS) [44], the Ecosystem Demography model (ED/ED2) [45,46], and Land Model 3 with PPA (LM3-PPA) [47], based on the Perfect Plasticity Approximation (PPA) [48,49]. These models represent the current state-of-the-art in modeling vegetation dynamics globally. While individual-based global models have begun to merge, forest landscape models have remained in between, focused on spatial processes of fire, harvest, and biological disturbance. Yet, previous research has shown that such forest landscape models are often insensitive to landscape configuration and are therefore aspatial [50], counter to the main assumption and selling point of these models.

While most forest landscape and terrestrial biosphere models lack individual trees, the SAS [45] and PPA [48,51,52] model reduction strategies have demonstrated an ability to successfully up-scale gap dynamics to forest stands. Other up-scaling strategies exist as well. One recent forest landscape model participating in the CoFoLaMo intercomparison scales from individual trees to stands by pre-computing light tables [53]. Regardless of the model structure, it is clear that gap, forest landscape, and terrestrial biosphere models are beginning to merge into new models of the terrestrial biosphere. This trend is also attributable to improvements in computational efficiency with new processor designs and cluster or cloud computing infrastructure. As few, if any, existing models are designed for highly parallel architectures (e.g., general-purpose graphics processing units, or GPGPUs), there remains much potential for future model efficiency gains.

In this forest biogeochemistry model intercomparison, we focus on two sites on the East Coast of the United States, Harvard Forest (HF), Massachusetts and Jones Ecological Research Center (JERC), Georgia. The two sites were selected for their representativeness of the United States Eastern Seaboard and for the availability of data needed to parameterize and validate the models. Harvard Forest is one of the most-studied forests in the world, with Google Scholar returning 12,700 results for the site. We focus on results for the Environmental Measurement Station (EMS) eddy covariance (EC) flux tower site within the Little Prospect Hill tract - the longest-running eddy covariance flux tower in the world. Previous research at the EMS EC flux tower site found unusually high rates of ecosystem respiration in winter and low rates in mid-to-late summer compared to other temperate forests [54]. While the mechanisms behind these observed patterns remains poorly understood, this observation is outside the scope of the presented research.

Between 1992 and 2004, the site acted as a net carbon sink, with a mean annual uptake rate of 2.5*MgCha*^−1^*year*^−1^. Aging dominated the site characteristics, with a 101-115 Mg C ha-1 increase in biomass, comprised predominantly of growth of red oak (*Quercus rubra*). The year 1998 showed a sharp decline in net ecosystem exchange (NEE) and other metrics, recovering thereafter [55]. As Urbanski *et al.* [55] note of the Integrated Biosphere Simulator 2 (IBIS2) and similar models at the time, “the drivers of interannual and decadal changes in NEE are long-term increases in tree biomass, successional change in forest composition, and disturbance events, processes not well represented in current models.” The two models used in the intercomparison study, a SORTIE-PPA [48,49] variant and LANDIS-II with NECN succession [56,57], are intended to directly address these model shortcomings.

While there have been fewer studies at Jones Ecological Research Center, Georgia, USA, Google Scholar returns 1,370 results for the site, reflecting its growing role in forest sciences research. Our study focuses on the Red Dirt (RD) EC flux tower site within the mesic sector, for which a handful of relevant studies exist. Two recent studies [58,59] indicate that the mesic sector of this subtropical pine savanna functions as a moderate carbon sink (NEE = −0.83 *Mg C ha*^−1^ *year*^−1^; −1.17 *Mg C ha*^−1^ *year*^−1^), reduced to near-neutral uptake during the 2011 drought (NEE = −0.17 *Mg C ha*^−1^*year*^−1^), and is a carbon source when prescribed burning is taken into account. NEE typically recovered to pre-fire rates within 30-60 days. The mechanisms behind soil respiration rates here again appear to be complex, site-specific, and poorly understood [59].

Overall, existing research highlights the importance of fire and drought to carbon exchange in long-leaf pine (*Pinus palustris*) and oak (*Quercus spp.*) savanna systems [58–60] at JERC. This is in contrast to the secondary growth-dominated deciduous broadleaf characteristics of Harvard Forest. Species diversity at the EMS tower site is 350% greater than that of the JERC-RD site, with 14 species from a variety of genera compared to four species from only two genera, *Pinus* and *Quercus*.

In this work, we aim to establish a foundation for future forest biogeochemistry model intercomparisons. This includes open-source object-oriented software to facilitate model parameterization, validation, intercomparison, and simplified reproducibility of results. We perform the model intercomparison for two key research forests in the United States to assess the ability of each model to reproduce observed biogeochemistry pools and fluxes over time. We hypothesize that the inclusion of forest growth, compositional change, and mortality processes in both models will allow for accurate predictions of biomass and NEE dynamics, as suggested in previous research Urbanski *et al.* [55]. Accordingly, we compare both models to observations and to each other for a host of metrics related to biomass, C, N, and forest composition at the two research sites.

## 2. Materials and Methods

LANDIS-II NECN and PPA-SiBGC were parameterized for two forested sites in the eastern United States, Harvard Forest, Massachusetts and Jones Ecological Research Center, Georgia. At the HF site, we focus on Little Prospect Hill and the EMS EC flux tower (HF-EMS). At the JERC site, we focus on the mesic zone and RD EC flux tower (JERC-RD). Both sites provided local EC and meteorological measurements to conduct this study. Plots of EC flux and meteorological tower measurements for both sites are located in Appendix A; maps of both sites are located in Appendix B.

Both models were parameterized using data available for each site, including local (i.e., field measurements) and general information sources (e.g., species compendiums and other published sources). As these empirical or observational values were used to parameterize both models, further model calibration (i.e., parameter tuning) was not necessary. This is because tuning parameters away from measured values to improve model performance, or defining a separate set of tuning parameters, is known to produce model over-fitting (i.e., reduced generality) and thus false improvements in model accuracy through reduced parsimony [61]. We explicitly avoided this practice, as it is only appropriate when fitting empirical growth-and-yield models such as Prognosis, also known as the Forest Vegetation Simulator (FVS) [18,19]. All model parameters are provided in the Appendix C. We close the methodology section with descriptions of the metrics, models, and criteria used in the intercomparisons.

### 2.1. Model Descriptions

In the following sections, we provide a brief overview of the two forest ecosystem models used in this intercomparison study. For detailed information on each model, readers are encouraged to refer to the original publications.

#### 2.1.1. LANDIS-II NECN

The LANDIS-II model is an extension of the original LANdscape DIsturbance and Succession (LANDIS) model [62–64] into a modular software framework [56]. Specifically, LANDIS-II is a model core containing basic state information that interfaces or communicates with external user-developed models known as “extensions” using a combination of object-oriented and modular design. This design makes LANDIS-II a modeling framework rather than a model. The LANDIS family of models, which also includes LANDIS PRO [65] and Fin-LANDIS [66,67], are stochastic hybrid models [40] based on the vital attributes/fuzzy systems approach of the LANDSIM model genre [68]. Perhaps unknowingly, this genre borrows heavily from cellular automata [69] and thus Markov Chains by applying simple heuristic rule-based systems, in the form of vital attributes, across two-dimensional grids.

Models of the LANDSIM genre focus on landscape-scale processes and assume game-theoretic vital attribute controls over successional trajectories following disturbance [70]. The LANDSIM model genre is thus a reasonable match for the classical forest fire model [71], given its local two-dimensional cellular basis. In contrast to the original LANDIS model, LANDIS-II is implemented in Microsoft C# rather than ISO C++98 [72], simplifying model development in exchange for a proprietary single-vendor software stack [56].

The latest version of LANDIS-II (v7) supports Linux through use of the Microsoft.NET Core developer platform. The modular design of LANDIS-II is intended to simplify the authorship and interaction of user-provided libraries for succession and disturbance. The centralized model core stores basic landscape and species state information and acts as an interface between succession and disturbance models. While there have been numerous forest landscape models over the years [21–25], the LANDIS family of models has enjoyed notable longevity and is currently united under the LANDIS-II Foundation. Part of its longevity is attributable to the prioritization of model functionality over realism in order to appeal to application-minded managers seeking a broad array of functionality.

The Net Ecosystem Carbon and Nitrogen (NECN) model [57] is a simplified variant of the classical Century model [34,35]. The original ten soil layers in Century have been replaced by a single soil layer, with functions for growth and decay borrowed directly from Century v4.5. The NECN succession model Figure 1 is thus a process-based model that simulates C and N dynamics along the plant-soil continuum at a native monthly timestep.

**Figure 1.**
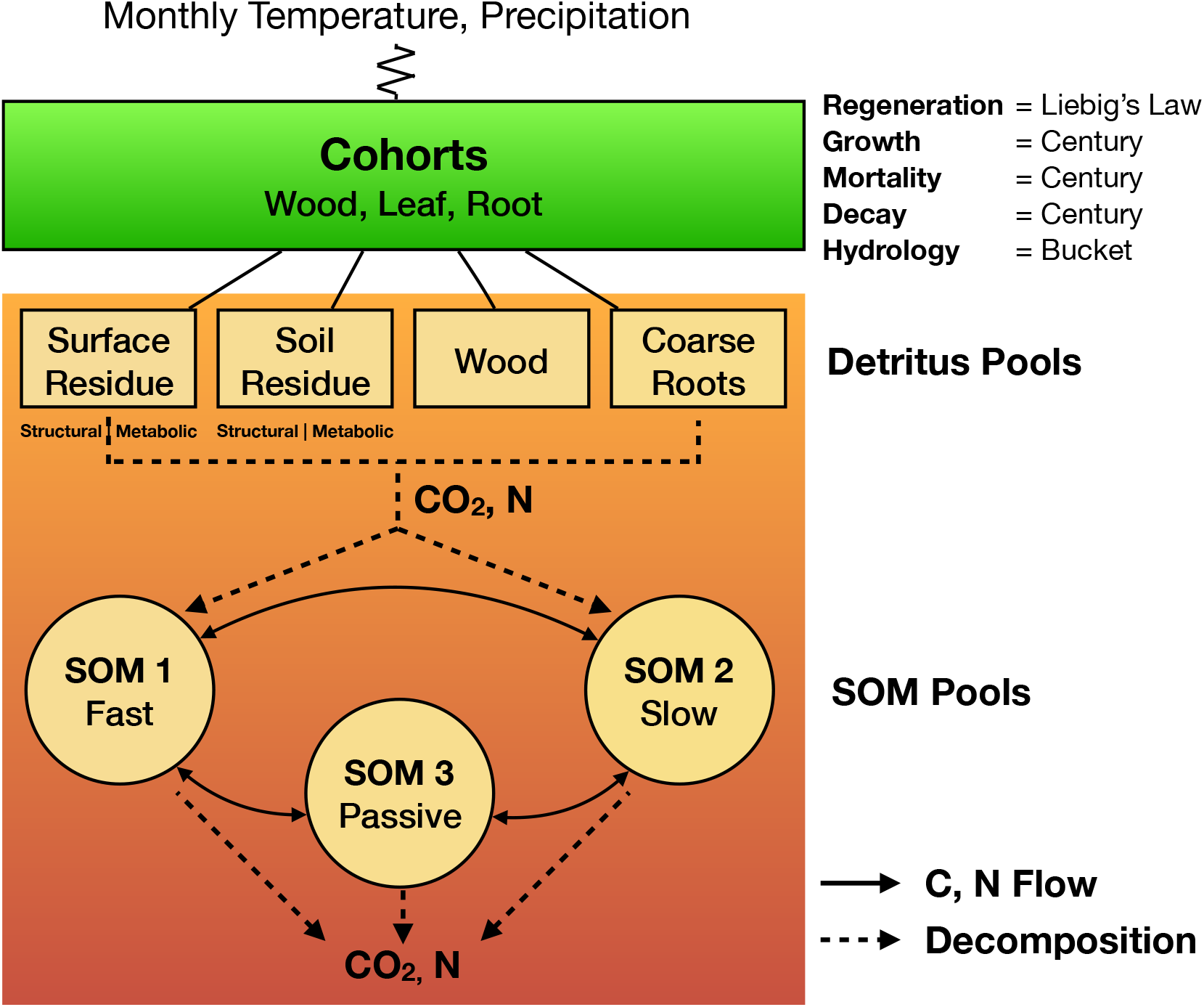
LANDIS-II NECN model structure

Atmospheric effects are included through monthly climate (i.e., temperature maxima, minima, means, and standard deviations, and precipitation means and standard deviations). Explicit geometric representation of tree canopies is forgone in favor of bounded statistical growth models based theoretically on Liebig’s Law of the Minimum. Functions for growth, mortality, and decay are adopted from Century [34] while hydrology is based on the simple bucket model [73]. The regeneration function is the only new process in NECN and is also based on Liebig’s Law. For a detailed description of the NECN model, readers may refer to the original model publication [57]. Parameterization of the LANDIS-II model for both sites was based on updating parameters used in recent [74–77] and ongoing (Flanagan et al., *in review*) work.

#### 2.1.2. PPA-SiBGC

The PPA-SiBGC model belongs to the SORTIE-PPA family of models [48,51] within the SAS-PPA model genre, based on a simple and analytically tractable approximation of the classical SORTIE gap model [78,79]. The Perfect Plasticity Approximation, or PPA [48,49], was derived from the dual assumptions of perfect crown plasticity (e.g., space-filling) and phototropism (e.g., stem-leaning), both of which were supported in empirical and modeling studies [51]. The discovery of the PPA was rooted in extensive observational and *in silico* research [48]. The PPA model was designed to overcome the most computationally challenging aspects of gap models in order to facilitate model scaling from the landscape to global scale.

The PPA and its predecessor, the size-and-age structured (SAS) equations [45,80], are popular model reduction techniques employed in current state-of-the-art terrestrial biosphere models [28]. The PPA model can be thought of metaphorically as Navier-Stokes equations of forest dynamics, capable of modeling individual tree population dynamics with a one-dimensional von Foerster partial differential equation [48]. The simple mathematical foundation of the PPA model is provided in Equation 1.

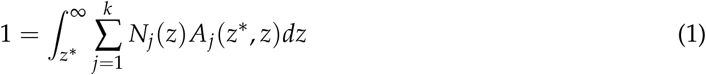

where *k* is the number of species, *j* is the species index, *N*_*j*_(*z*) is the density of species *j* at height *z, A*_*j*_(*a*\*, *z*) is the projected crown area of species *j* at height *z*, and *dz* is the derivative of height. In other words, we discard the spatial location of individual trees and calculate the height at which the integral of tree crown area is equal to the ground area of the stand. This height is known as the theoretical *z*\* height, which segments trees into overstory and understory classes [48].

The segmentation of the forest canopy into understory and overstory layers allows for separate coefficients or functions for growth, mortality, and fecundity to be applied across strata, whose first moment accurately approximates the dynamics of individual-based forest models. Recent studies have shown that the PPA model faithfully reduces the dynamics of the more recent neighborhood dynamics (ND) SORTIE-ND gap model [81] and is capable of accurately capturing forest dynamics [82,83].

In this work, we applied a simple biogeochemistry variant of the SORTIE-PPA model, PPA-SiBGC [Erickson and Strigul, *In Review*] Figure 2.

**Figure 2.**
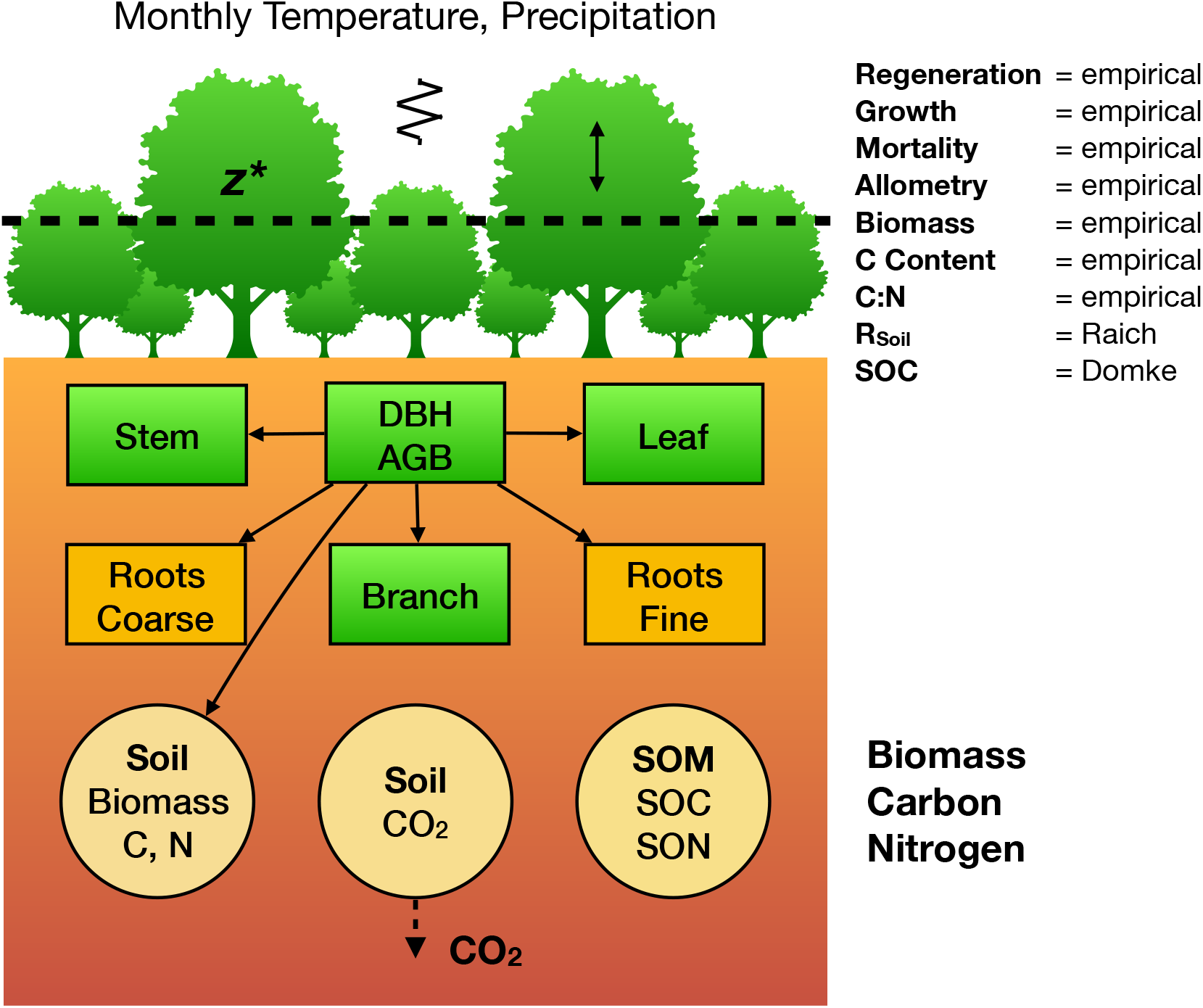
PPA-SiBGC model structure; Raich *et al.* [84]; Domke *et al.* [85]

Empirical observations were relied upon for the C and N content of tree species compartments. Stoichiometric relations were used to estimate N from C, based on empirical measurements provided for both sites. All values were calculated directly from observations. Previously published equations [86] and parameters [87] were used to model crown allometry. Together with inventory data, general biomass equations were used to estimated dry weight mass (*kg*) for tree stems, branches, leaves, and, fine and coarse roots [88]. Carbon content is assumed to be 50% of dry mass, supported by data. Monthly soil respiration is modeled using the approach of Raich *et al.* [84], while soil organic C is modeled using the simple generalized approach of Domke *et al.* [85]. Species- and stratum-specific parameters for growth, mortality, and fecundity were calculated from observational data available for both sites.

### 2.2. Site Descriptions

In the following sections, we describe the two forested sites on the East Coast of the United States: HF-EMS and the JERC-RD. A critical factor in the selection of the sites was the availability of eddy covariance flux tower data needed to validate NEE in the models.

#### 2.2.1. HF-EMS

The HF-EMS EC flux tower is located within the Little Prospect Hill tract of Harvard Forest (42.538° N, 72.171° W, 340 m elevation) in Petersham, Massachusetts, approximately 100 km from the city of Boston [55]. The tower has been recording NEE, heat, and meteorological measurements since 1989, with continuous measurements since 1991, making it the longest-running eddy covariance measurement system in the world. The site is currently predominantly deciduous broadleaf second-growth forests approximately 75-95 years in age, based on previous estimates [89]. Soils at Harvard Forest originate from sandy loam glacial till and are reported to be mildly acidic [55].

The site is dominated by red oak (*Quercus rubra*) and red maple (*Acer rubrum*) stands, with sporadic stands of Eastern hemlock (*Tsuga canadensis*), white pine (*Pinus strobus*), and red pine (*Pinus resinosa*). When the site was established, it contained 100 Mg C ha^−1^ in live aboveground woody biomass [89]. As noted by Urbanski *et al.* [55], approximately 33% of red oak stands were established prior to 1895, 33% prior to 1930, and 33% before 1940. A relatively hilly and undisturbed forest (since the 1930s) extends continuously for several km^2^ around the tower. In 2000, harvest operations removed 22.5 Mg C ha^−1^ of live aboveground woody biomass about 300 m S-SE from the tower, with little known effect on the flux tower measurements. The 40 biometric plots were designated via stratified random sampling within eight 500 m transects Urbanski *et al.* [55]. The HF-EMS tower site currently contains 34 biometric plots at 10 m radius each, covering 10,681 m^2^, or approximately one hectare, in area. Summary statistics for the EMS tower site for the year 2002 are provided in Table 1.

**Table 1.** HF-EMS forest inventory summary for the 34 tower plots in 2002; DBH = depth at breast height (cm); BA = basal area per hectare (m^2^); Stocking = n*_trees_* per hectare; QMD = quadratic mean diameter (cm); SDI = Reineke’s stand density index [90]

| Statistic | N | Mean | St. Dev. | Min | Pctl(25) | Pctl(75) | Max |
| --- | --- | --- | --- | --- | --- | --- | --- |
| DBH<sub>Mean</sub> | 34 | 24.64 | 3.50 | 17.32 | 22.39 | 27.13 | 31.97 |
| DBH<sub>St.Dev.</sub> | 34 | 10.92 | 2.74 | 6.11 | 8.60 | 12.51 | 16.88 |
| BA | 34 | 35.29 | 11.55 | 13.78 | 25.98 | 42.34 | 57.31 |
| Stocking | 34 | 639.43 | 232.38 | 318.31 | 421.76 | 787.82 | 1,145.92 |
| QMD | 34 | 26.87 | 4.00 | 18.34 | 23.91 | 29.64 | 34.39 |
| SDI | 34 | 680.13 | 214.45 | 280.56 | 531.35 | 853.97 | 1,071.37 |

A table of observed species abundances for the year 2002 are provided in Table 2, using tree species codes from the USDA PLANTS database (https://plants.usda.gov).

**Table 2.** HF-EMS species abundance for the 34 tower plots in 2002

| Species | Count |
| --- | --- |
| ACPE | 56 |
| ACRU | 4,924 |
| BEAL | 729 |
| BELE | 239 |
| BEPO | 116 |
| CADE | 1 |
| FAGR | 277 |
| FRAM | 258 |
| ILVE | 86 |
| PIGL | 397 |
| PIRE | 638 |
| PIST | 582 |
| PRSE | 270 |
| QURU | 2,485 |
| QUVE | 247 |
| TSCA | 1,926 |

Data were collected here for a range of studies, as evidenced by the Harvard Forest Data Archive. Datasets used in model validation include HF001-04, HF004-02, HF069-09, HF278-04, HF069-06, HF015-05, HF006-01, and HF069-13. These include weather station and forest inventory time-series, eddy covariance flux tower measurements, soil respiration, soil organic matter, and studies on C:N stoichiometry. Standard measurement techniques were used for each. For both sites, local tree species, age, depth-at-breast-height (DBH), biomass, soil, and meteorological data were primarily used to parameterize the models.

#### 2.2.2. JERC-RD

Jones Ecological Research Center at Ichauway is located near Newton, Georgina, USA (31° N, 84° W, 25-200 m elevation). The site falls within the East Gulf Coastal Plain and consists of flat to rolling land sloping to the southwest. The region is characterized by a humid subtropical climate with temperatures ranging from 5-34 °C and precipitation averaging 132 cm year-1. The overall site is 12,000 ha in area, 7,500 ha of which are forested [91]. The site also exists within a tributary drainage basin that eventually empties into the Flint River. Soils here are underlain by karst Ocala limestone and mostly Typic Quartzipsamments, with sporadic Grossarenic and Aquic Arenic Paleudults [92]. Soils here often lack well-developed organic horizons [91–93].

Forests here are mostly second-growth, approximately 65-95 years in age. Long-leaf pine (*Pinus palustris*) dominates the overstory, while the understory is comprised primarily of wiregrass (*Aristida stricta*) and secondarily of shrubs, legumes, forbs, immature hardwoods, and regenerating long-leaf pine forests [94]. Prescribed fire is a regular component of management here, with stands often burned at regular 1-5 year intervals [91]. This has promoted wiregrass and legumes in the understory, while reducing the number of hardwoods [91]. The RD EC flux tower is contained within the mesic/intermediate sector. This site consists of only four primary tree species from two genera: long-leaf pine (*Pinus palustris*), water oak (*Quercus nigra*), southern live oak (*Quercus virginiana*), and bluejack oak (*Quercus incana*). Measurements for the RD tower are available for the 2008-2013 time period. Summary statistics for the RD tower site for the year 2008 are provided in Table 3.

**Table 3.**
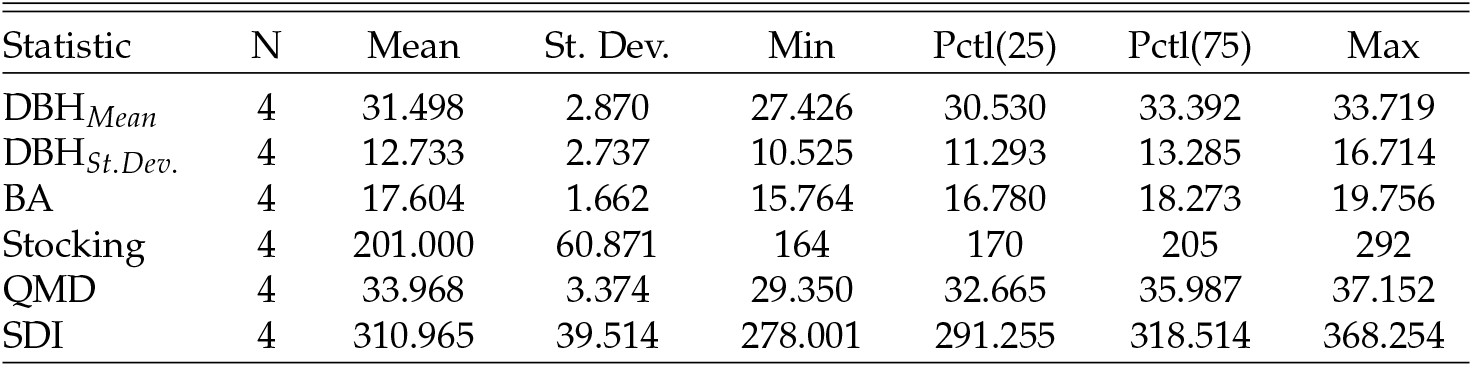
JERC-RD forest inventory summary for the 4 tower plots in 2009; DBH = depth at breast height (cm); BA = basal area per hectare (m^2^); Stocking = n*_trees_* per hectare; QMD = quadratic mean diameter (cm); SDI = Reineke’s stand density index [90]

A table of observed species abundances for the year 2009 are provided in Table 4.

**Table 4.** JERC-RD species abundance for the 4 tower plots in 2009

| Species | Count |
| --- | --- |
| PIPA | 2,063 |
| QUIN | 14 |
| QUNI | 22 |
| QUVI | 66 |

Datasets used in model validation at JERC-RD include JC010-02, JC010-01, JC003-04, JC004-01, JC003-07, and JC011-01. These include weather station and eddy covariance flux tower measurements, forest inventory data, soil respiration, soil organic matter, and studies on C:N stoichiometry. Standard measurement techniques were also used for each of these.

### 2.3. Site Data

To conduct this model intercomparison exercise at HF-EMS, we leveraged the large amount of data openly available to the public through the Harvard Forest Data Archive:

> http://harvardforest.fas.harvard.edu/harvard-forest-data-archive

Jones Ecological Research Center has hosted multiple research efforts over the years, collectively resulting in the collection of a large data library. However, JERC-RD site data are not made openly available to the public and are thus only available by request. One may find contact information located within their website:

> http://www.jonesctr.org

### 2.4. Scales, Metrics, and Units

The selection of simulation years was based on the availability of EC flux tower data used in model validation. Thus, we simulated the HF-EMS site for the years 2002-2012 and the JERC-RD site for the years 2009-2013. For both sites and models, we initialized the model state in the first year of simulations using field observations. The PPA-SiBGC model used an annual timestep while LANDIS-II NECN used a monthly timestep internally. Both models may be set to other timesteps if desired.

The areal extent of the single-site model intercomparisons were designed to correspond to available field measurements. At both sites, tree inventories were conducted in 10,000 m^2^, or one-hectare, areas. All target metrics were converted to an annual areal basis to ease interpretation, comparison, and transferability of results. Importantly, an areal conversion will allow comparison to other sites around the world. While flux tower measurements for both sites were already provided on an areal (m^−2^) basis, many other variables were converted to harmonize metrics between models and study sites. For example, moles CO_2_ measurements were converted to moles C through well-described molecular weights, all other measures of mass were converted to kg, and all areal and flux measurements were harmonized to m^−2^. A table of metrics and units used in the intercomparison of LANDIS-II and PPA-SiBGC is provided in Table 5.

**Table 5.** Model intercomparison abbreviations, metrics, and units

| Abbreviation | Metric | Units |
| --- | --- | --- |
| NEE | Net ecosystem exchange | kg C m<sup>-2</sup> year<sup>-1</sup> |
| B<sub>AG</sub> | Aboveground biomass | kg mass m<sup>-2</sup> |
| C<sub>AG</sub> | Aboveground C | kg C m<sup>-2</sup> |
| N<sub>AG</sub> | Aboveground N | kg N m<sup>-2</sup> |
| B<sub>BG</sub> | Belowground biomass | kg mass m<sup>-2</sup> |
| C<sub>BG</sub> | Belowground C | kg C m<sup>-2</sup> |
| N<sub>BG</sub> | Belowground N | kg N m<sup>-2</sup> |
| C<sub>SO</sub> | Soil organic C | kg C m<sup>-2</sup> |
| N<sub>SO</sub> | Soil organic N | kg N m<sup>-2</sup> |
| r<sub>soil</sub> | Soil respiration C | kg C m<sup>-2</sup> year<sup>-1</sup> |
| ANPP | Aboveground net primary production | kg mass m<sup>-2</sup> year<sup>-1</sup> |
| B<sub>Sp</sub> | Species aboveground biomass | kg mass m<sup>-2</sup> |
| n<sub>Sp</sub> | Species relative abundance | % |

In the subsequent section, we describe the model intercomparison methodology.

### 2.5. Model Intercomparison

Intercomparison of the PPA-SiBGC and LANDIS-II models at the HF-EMS and JERC-RD EC flux tower sites was conducted using a collection of object-oriented functional programming scripts written in the R language for statistical computing [95]. These scripts were designed to simplify model configuration, parameterization, operation, calibration/validation, plotting, and error calculation. The scripts and our parameters are available on GitHub (https://github.com/adam-erickson/ecosystem-model-comparison), making our results fully and efficiently reproducible. The R scripts are also designed to automatically load and parse the results from previous model simulations, in order to avoid reproducibility issues stemming from model stochasticity. We use standard regression metrics applied to the time-series of observation and simulation data to assess model fitness. The metrics used include the coefficient of determination (*R*^2^), root mean squared error (RMSE), mean absolute error (MAE), and mean error (ME) or bias, calculated using simulated and observed values. Our implementation of *R*^2^ follows the Bravais-Pearson interpretation as the squared correlation coefficient between observed and predicted values [96]. This implementation is provided in Equation 2.

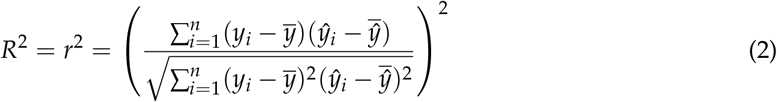

where *n* is the sample size, *y*_*i*_ is the *i*th observed value, *ŷ*_*i*_ is the *i*th predicted value, 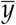 is the mean observed value, and 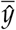 is the mean predicted value. The calculation of RMSE follows the standard formulation, as shown in Equation 3.

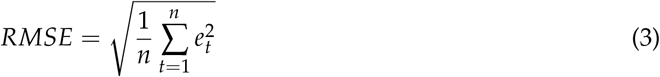

where *n* is the sample size and *e*_*t*_ is the error for the *t*th value, or the difference between observed and predicted values. The calculation of MAE is similarly unexceptional, per Equation 4.

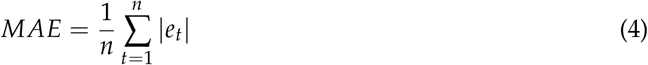

where again *n* is the sample size and *e*_*t*_ is the error for the *t*th value. Our calculation of mean error (ME) or bias is the same as MAE, but without taking the absolute value.

While Nash-Sutcliffe efficiency (NSE) is often used in a simulation model context, we selected the Bravais-Pearson interpretation of *R*^2^ over NSE to simplify the interpretation of results. The NSE metric replaces 1 − (*SS*_*predictions*_/*SS*_*observations*_) with (*SS*_*observations*_ *− SS*_*predictions*_)/*SS*_*observations*_, where *SS* is the sum of squares. Thus, NSE is analogous to the standard *R*^2^ coefficient of determination used in regression analysis [97]. The implementation of *R*^2^ that we selected is important to note, as its results are purely correlative and quantify only dispersion, ranging in value between 0 and 1. This has some desirable properties in that no negative or large values are produced, and that it is insensitive to differences in scale. Regardless of the correlation metric used, complementary metrics are needed to quantify the direction (i.e., bias) and/or magnitude of error. We rely on RMSE and MAE to provide information on error or residual magnitude, and ME to provide information on bias. We utilize a visual analysis to assess error directionality over time, as this can be poorly characterized by a single coefficient, masking periodicity.

We compute *R*^2^, RMSE, MAE, and ME for time-series of the metrics described in Table 5 on page 11. These include NEE, above- and below-ground biomass, C, and N, soil organic C and N, soil respiration (*r*_*soil*_), aboveground net primary production (ANPP), and, species aboveground biomass and relative abundance. All of these metrics are pools with the exception of NEE, *r*_*soil*_, and ANPP fluxes. Finally, we diagnose the ability of both models to meet a range of logistical criteria related to deployment: *model usability, performance, and transferability*. Model usability is assessed per four criteria:

1. Ease of installation
2. Ease of parameterization
3. Ease of program operation
4. Ease of parsing outputs

Model software performance is assessed per a single metric: the speed of program execution for each site for the predefined simulation duration. The durations are 11 years and 5 years for the HF-EMS and JERC-RD EC flux tower sites, respectively. Simulation results are output at annual temporal resolution, the standard resolution for both models; while NECN operates on a monthly timestep, most other modules of LANDIS-II are annual. Finally, model transferability is assessed per the following five criteria:

1. Model generalizability
2. Availability of parameterization data
3. Size of the program
4. Cross-platform support
5. Ease of training new users

Each of these logistical criteria are compared in a qualitative analysis, with the exception of software performance.

## 3. Results

Both PPA-SiBGC and LANDIS-II NECN showed strong performance for pools at the two model intercomparison sites, frequently achieving *R*^2^ values approaching unity. Yet, both models showed weak performance for fluxes. The models failed to accurately predict ANPP, while PPA-SiBGC showed stronger *r*_*soil*_ performance and LANDIS-II NECN showed stronger NEE performance. The *R*^2^ values for both models and sites are visualized in Figure 3.

**Figure 3.**
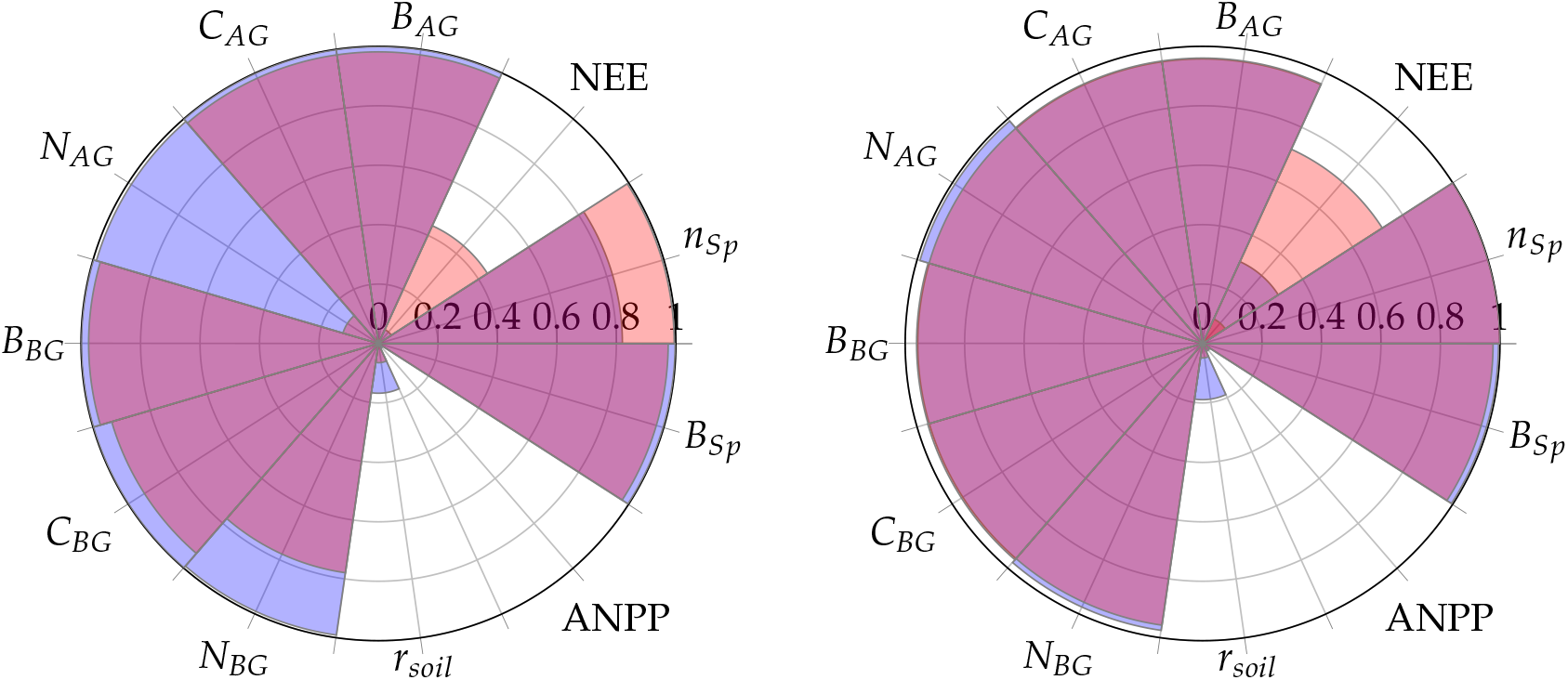
Overall model performance (R^2^) for both models and sites; left = HF-EMS; right = JERC-RD; periwinkle = PPA-SiBGC; pink = LANDIS-II NECN; violet = intersection

On average, PPA-SiBGC outperformed LANDIS-II NECN across the sites and metrics tested, showing higher correlations, lower error, and less bias overall (HF-EMS 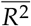 = 0.73, +0.07, 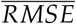 = 4.84, −0.39, 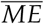 = −1.18, −3.70; JERC-RD 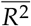 = 0.76, +0.04, 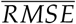 = 2.69, −0.17, 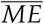 = 0.78, +0.53). This result is based on calculating mean values for *R*^2^, RMSE, MAE, and ME in order to clearly translate the overall results. The two models produced the following mean values for each of the four statistical metrics and two sites:

**Table 6.** Overall mean values across each of the sites and metrics tested

| Metric | PPA-SiBGC |  |  |  | LANDIS-II NECN |  |  |  |
| --- | --- | --- | --- | --- | --- | --- | --- | --- |
| | $R^2$ | RMSE | MAE | ME | $R^2$ | RMSE | MAE | ME |
| Mean | 0.74 | 3.77 | 3.58 | -0.20 | 0.69 | 9.60 | 8.73 | 2.31 |

As shown in Table 6, PPA-SiBGC yielded higher *R*^2^ values and lower RMSE, MAE, and ME values in comparison to LANDIS-II, on average, across all sites and metrics tested. Below, we provide model intercomparison results individually for the two sites, HF-EMS and JERC-RD.

### 3.1. HF-EMS

For the HF-EMS site, PPA-SiBGC showed higher *R*^2^ values and lower RMSE, MAE, and ME values compared to LANDIS-II NECN across the range of metrics. While PPA-SiBGC predicted NEE and species relative abundance showed weaker correlations with observed values compared to LANDIS-II NECN, the magnitude of error was lower, as evidenced by lower RMSE, MAE, and ME values. While LANDIS-II NECN showed a lower magnitude of error for belowground N, this is the only metric where this is the case, while the correlation of this metric to observed values was also lower than that of PPA-SiBGC. Overall results for the HF-EMS site model intercomparison are shown in Table 7.

**Table 7.** Model fitness for HF-EMS

| Metric | PPA-SiBGC |  |  |  | LANDIS-II NECN |  |  |  |
| --- | --- | --- | --- | --- | --- | --- | --- | --- |
| | $R^2$ | RMSE | MAE | ME | $R^2$ | RMSE | MAE | ME |
| NEE | 0.05 | 0.19 | 0.16 | -0.03 | 0.44 | 0.49 | 0.44 | 0.44 |
| $B_{AG}$ | 1.00 | 10.12 | 10.11 | 10.11 | 0.98 | 2.48 | 2.48 | -2.48 |
| $C_{AG}$ | 1.00 | 0.03 | 0.03 | -0.02 | 0.98 | 1.24 | 1.24 | -1.24 |
| $N_{AG}$ | 0.99 | 1.44 | 1.44 | -1.44 | 0.12 | 1.99 | 1.99 | -1.99 |
| $B_{BG}$ | 1.00 | 9.09 | 9.08 | 9.08 | 0.97 | 2.82 | 2.82 | -2.82 |
| $C_{BG}$ | 1.00 | 7.82 | 7.81 | -7.81 | 0.93 | 9.87 | 9.86 | -9.86 |
| $N_{BG}$ | 0.99 | 0.56 | 0.56 | 0.56 | 0.78 | 0.12 | 0.12 | -0.12 |
| $r_{soil}$ | 0.17 | 0.63 | 0.62 | -0.62 | 0.06 | 1.10 | 1.10 | -1.10 |
| ANPP | 0.02 | 0.20 | 0.20 | -0.20 | 0.0002 | 0.82 | 0.79 | 0.73 |
| $C_{SO}$ | ... | 26.49 | 26.49 | -26.49 | ... | 36.63 | 36.63 | -36.63 |
| $N_{SO}$ | ... | 1.33 | 1.33 | -1.33 | ... | 1.60 | 1.60 | -1.60 |
| $B_{Sp}$ | 1.00 | 5.02 | 2.89 | 2.89 | 0.97 | 133.70 | 119.87 | 119.87 |
| $n_{Sp}$ | 0.82 | 0.05 | 0.03 | 0 | 0.99 | 0.29 | 0.22 | 0.22 |
| Mean | 0.73 | 4.84 | 4.67 | -1.18 | 0.66 | 14.86 | 13.78 | 4.88 |

Time-series figures allow a visual analysis of the temporal dynamics between observations and model predictions in order to assess the ability of models to capture interannual variability. Both models effectively captured temporal dynamics in biomass, C, and, species biomass and abundance. In Figure 4, the temporal differences in modeled NEE and aboveground C are shown for the two models in comparison to observations for the HF-EMS site. While LANDIS-II NECN predicted NEE showed a higher correlation with observations, the magnitude of error and bias were also higher. Furthermore, LANDIS-II NECN predicted that the HF-EMS site is a net C source, rather than sink, in contrary to observations. Meanwhile, PPA-SiBGC outperformed LANDIS-II NECN in aboveground C per both *R*^2^ and RMSE. Both models overpredicted species cohort biomass, while LANDIS-II NECN underpredicted total aboveground C.

**Figure 4.**
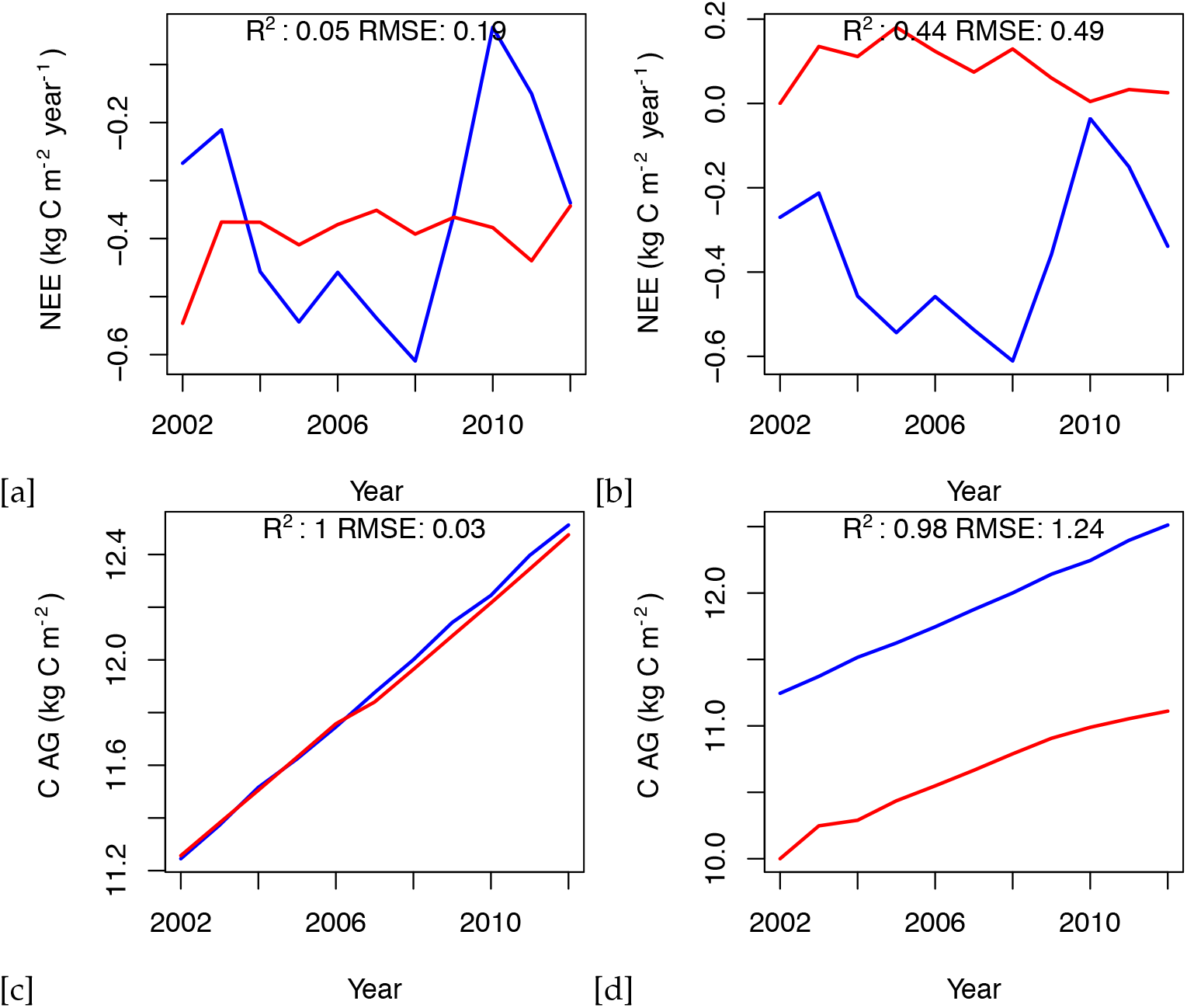
Simulated and observed NEE and aboveground C for the HF-EMS site; observations = blue; simulations = red; a = PPA-SiBGC NEE; b = LANDIS-II NECN NEE; c = PPA-SiBGC *C*_*AG*_; d = LANDIS-II NECN *C*_*AG*_

An analysis of simulated species biomass and abundance also shows greater fidelity of the PPA-SiBGC model to data, as shown in Figure 5. As LANDIS-II NECN does not contain data on individual trees, species relative abundance is calculated based on the number of cohorts of each species. Two species were simulated in LANDIS-II NECN, as there are no explicit trees in the model and the number of cohorts appears to have no effect on the total biomass. Results for PPA-SiBGC indicate that species relative abundance may be improved in future studies by optimizing mortality and fecundity rates. Meanwhile, species biomass predictions output by LANDIS-II NECN were inverted from those of the observations.

**Figure 5.**
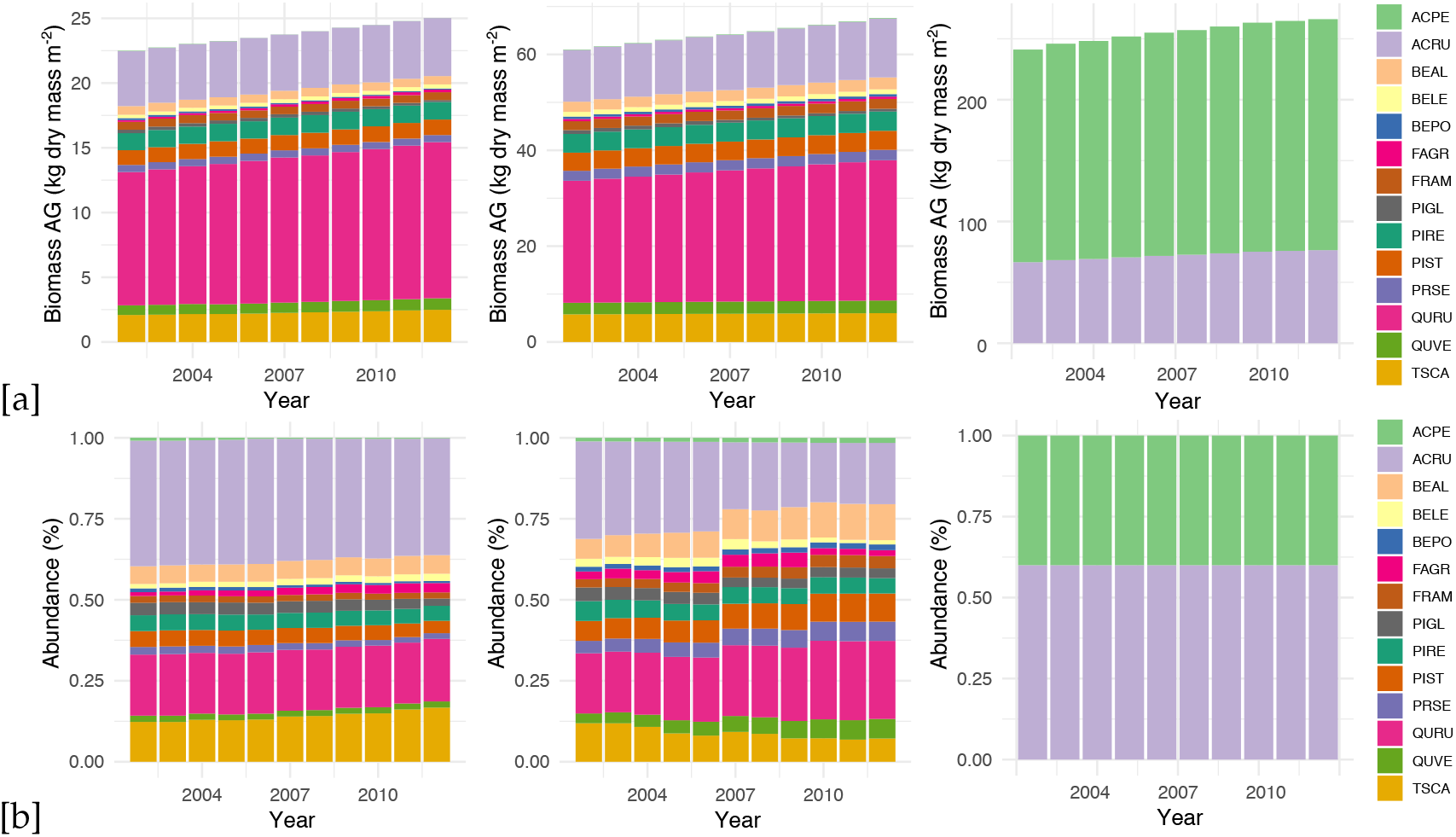
HF-EMS: Simulated and observed species aboveground biomass and relative abundance; a = biomass; b = abundance; left = observations, middle = PPA-SiBGC, right = LANDIS-II NECN; note that different scales are used for biomass

### 3.2. JERC-RD

For the JERC-RD site, both models showed stronger fidelity to data than for the HF-EMS site. Again, PPA-SiBGC showed higher *R*^2^ values and lower RMSE and MAE values compared to LANDIS-II NECN across the range of metrics tested. Yet, the margin between models was smaller for the JERC RD site. While PPA-SiBGC demonstrated higher correlations and lower errors for most metrics tested, LANDIS-II NECN outperformed PPA-SiBGC in a few cases. This includes lower error magnitude for NEE, aboveground N, belowground biomass, SOC, and SON. However, PPA-SiBGC showed correlations equal or higher for all metrics tested, and lower errors for all other metrics. Overall results for the JERC-RD site model intercomparison are shown in Table 8.

**Table 8.** Model fitness for JERC-RD

| Metric | PPA-SiBGC |  |  |  | LANDIS-II NECN |  |  |  |
| --- | --- | --- | --- | --- | --- | --- | --- | --- |
| | $R^2$ | RMSE | MAE | ME | $R^2$ | RMSE | MAE | ME |
| NEE | 0.30 | 1.68 | 1.64 | -1.64 | 0.09 | 0.13 | 0.11 | -0.05 |
| $B_{AG}$ | 0.96 | 1.48 | 1.47 | 1.47 | 0.96 | 9.77 | 9.76 | -9.76 |
| $C_{AG}$ | 0.96 | 1.63 | 1.63 | -1.63 | 0.96 | 4.88 | 4.88 | -4.88 |
| $N_{AG}$ | 0.99 | 0.29 | 0.29 | 0.29 | 0.96 | 0.05 | 0.05 | -0.05 |
| $B_{BG}$ | 0.96 | 10.84 | 10.83 | 10.83 | 0.96 | 1.37 | 1.20 | 1.20 |
| $C_{BG}$ | 0.96 | 5.26 | 5.26 | -5.26 | 0.96 | 6.46 | 6.46 | -6.46 |
| $N_{BG}$ | 0.98 | 1.44 | 1.44 | -1.44 | 0.96 | 1.60 | 1.60 | -1.60 |
| $r_{soil}$ | 0.19 | 0.78 | 0.71 | -0.67 | 0.05 | 2.40 | 2.40 | -2.40 |
| ANPP | 0.03 | 0.39 | 0.37 | -0.37 | 0.03 | 0.48 | 0.46 | 0.25 |
| $C_{SO}$ | ... | 4.30 | 4.30 | 4.30 | ... | 0.17 | 0.17 | -0.17 |
| $N_{SO}$ | ... | 0.38 | 0.38 | 0.38 | ... | 0.12 | 0.12 | 0.12 |
| $B_{Sp}$ | 1.00 | 6.47 | 3.90 | 3.90 | 0.98 | 28.97 | 20.52 | 20.52 |
| $n_{Sp}$ | 1.00 | 0.02 | 0.01 | -0 | 1.00 | 0.09 | 0.09 | -0.02 |
| Mean | 0.76 | 2.69 | 2.48 | 0.78 | 0.72 | 4.34 | 3.68 | -0.25 |

While both models showed higher performance at the JERC-RD site, an analysis of simulated species biomass and abundance again indicates greater fidelity of the PPA-SiBGC model to data, as shown in Figure 6. While LANDIS-II NECN overpredicts the rate of longleaf pine growth, PPA-SiBGC nearly perfectly matches observed species abundance and biomass trajectories for all species present. While the correlations are high, PPA-SiBGC overpredicts the magnitude of biomass here.

**Figure 6.**
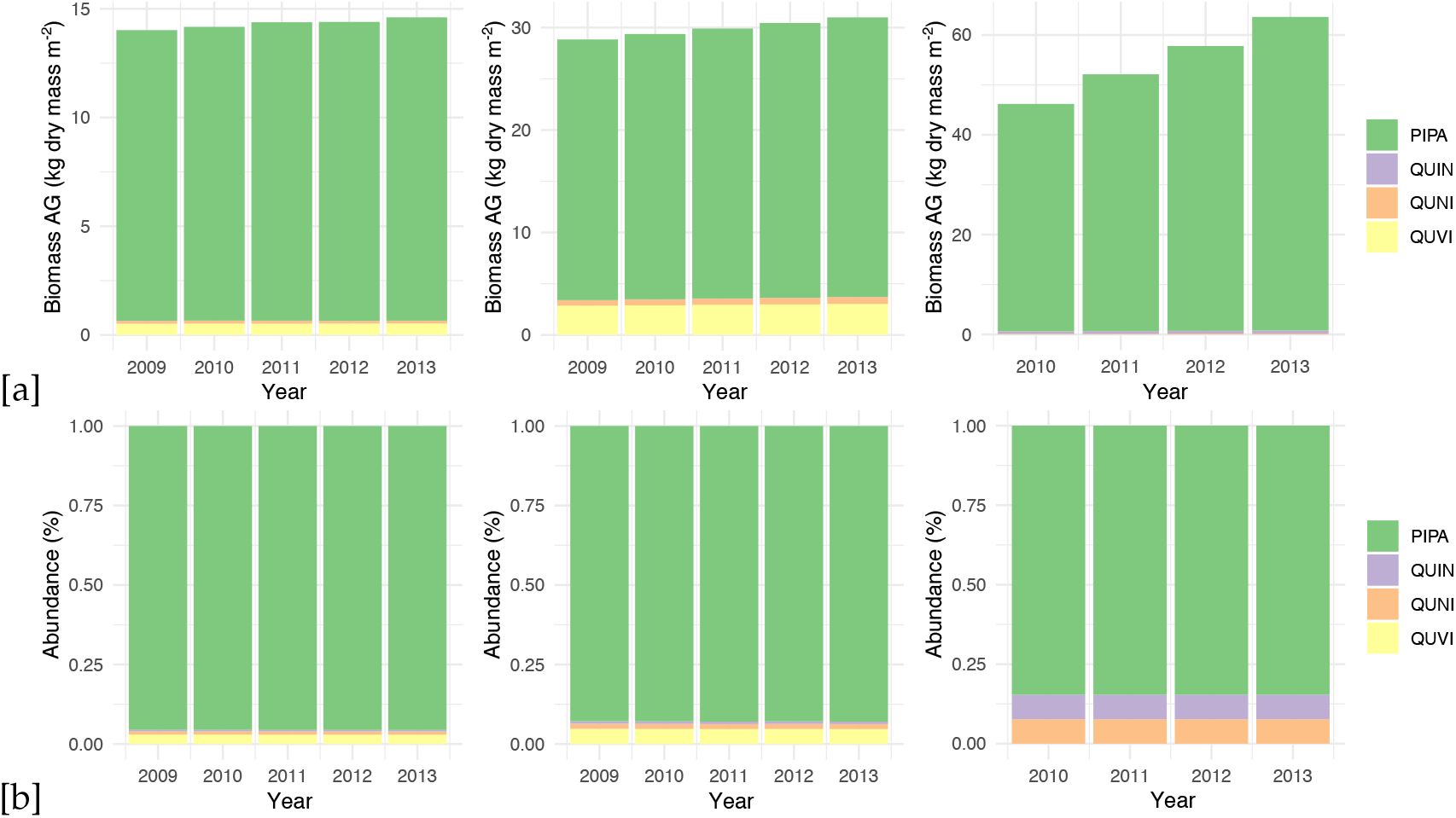
JERC-RD: Simulated and observed species aboveground biomass and relative abundance; a = biomass; b = abundance; left = observations, middle = PPA-SiBGC, right = LANDIS-II NECN; note that different scales are used for biomass

Our results for the HF-EMS and JERC-RD site model intercomparison exercise clearly indicate strong performance for both models at both sites. Results for the JERC-RD site are particularly close between the two models. Next, we assess results related to the logistics of model deployment to new computers, users, and modeling sites.

### 3.3. Model Usability, Performance, and Transferability

While the two models share a similar basis in forest dynamics and biogeochemistry modeling, they differ in important practical and conceptual terms. The command-line version of the PPA-SiBGC model used in this work, version 5.0, consists of approximately 500 lines of R code and is thus readily cross-platform, including cloud providers. Meanwhile, the LANDIS-II model core and NECN succession extension are an estimated 2,000 and 0.5 million lines of code, respectively. While this version of PPA-SiBGC fuses an explicit tree canopy geometry model with empirical data on fecundity, growth, mortality, and stoichiometry, the NECN extension of LANDIS-II borrows heavily from the process-based Century model [35], similar to the MAPSS-Century-1 (MC1) model [98]. This carries important implications for model parameterization needs. While PPA-SiBGC relies on typical forest inventory data, including tree species, age/size, and densities, LANDIS-II relies on species age/size and traits in the form of vital attributes, in addition to NECN parameters. Below, we summarize our findings regarding the logistics of model deployment.

#### 3.3.1. Model Usability

In the following section, we provide an assessment of model usability based on four criteria.

1. *Ease of installation* While LANDIS-II NECN requires the installation of two Windows programs, depending on the options desired, PPA-SiBGC is contained in a single R script and requires only a working R installation.
2. *Ease of parameterization* While both models can be difficult to parameterize for regions with little to no observational data, the simple biogeochemistry in PPA-SiBGC requires an order of magnitude fewer parameters than LANDIS-II NECN. In addition, PPA-SiBGC uses commonly available forest inventory data while NECN requires a number of parameters that may be difficult to locate.
3. *Ease of program operation* Both models use a command-line interface and are thus equally easy to operate. Yet, PPA-SiBGC is cross-platform and uses comma-separated-value (CSV) files for input tables, which are easier to work with than multiple tables nested within an unstructured text files. This additionally allows for simplification in designing model application programming interfaces (APIs), or model wrappers, a layer of abstraction above the models. These abstractions are important for simplifying model operation and reproducibility, and enable a number of research applications.
4. *Ease of parsing outputs* All PPA-SiBGC outputs are provided in CSV files in a single folder while LANDIS-II NECN generates outputs in multiple formats in multiple folders. While the PPA-SiBGC format is simpler and easier to parse, the image output formats used by LANDIS-II carry considerable benefit for spatial applications. Both models may benefit by transitioning spatiotemporal data to the NetCDF scientific file format used by most general circulation and terrestrial biosphere models.

#### 3.3.2. Model Performance

Next, we assess model performance in terms of the speed of operation on a consumer-off-the-shelf (COTS) laptop computer with a dual-core 2.8 GHz Intel Core i7-7600U CPU and 16 GB of DDR4-2400 RAM. We focus on a single performance metric, the timing of simulations. Other aspects of model performance in the form of precision and accuracy are described in previous sections. As shown in Table 9, PPA-SiBGC was between 1,200 and 2,800% faster than LANDIS-II NECN in our timing tests. This was surprising given that PPA-SiBGC models true cohorts (i.e., individual trees) in an interpreted language while LANDIS-II models theoretical cohorts (i.e., cohorts without a physical basis) in a compiled language. The difference in speed is likely attributable to the parsimony of the PPA-SiBGC model.

**Table 9.**
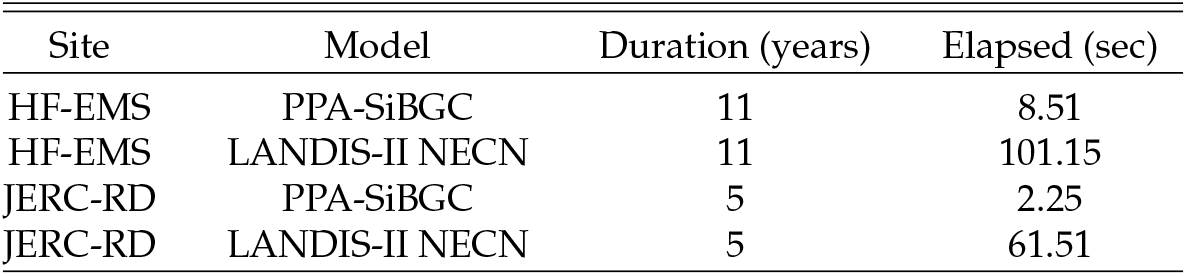
Simulation timing results

| Site | Model | Duration (years) | Elapsed (sec) |
| --- | --- | --- | --- |
| HF-EMS | PPA-SiBGC | 11 | 8.51 |
| HF-EMS | LANDIS-II NECN | 11 | 101.15 |
| JERC-RD | PPA-SiBGC | 5 | 2.25 |
| JERC-RD | LANDIS-II NECN | 5 | 61.51 |

#### 3.3.3. Model Transferability

Here, we discuss model transferability. In this section, we assess the effort required to transfer the models to new locations, new computer systems, or new users. All three are important logistical criteria for effective model deployment.

1. *Model generalization* Both models appear to generalize effectively to different forested regions of the world, as both have shown strong performance in this study and others. No clear winner is evident in this regard. In terms of model realism, PPA-SiBGC has a more realistic representation of forest canopies while LANDIS-II NECN has more realistic processes, as it is a Century model variant.
2. *Availability of parameterization data* While LANDIS-II NECN requires substantially greater parameterization data compared to PPA-SiBGC, it may often be possible to rely on previously published parameters. Meanwhile, the growth, mortality, and fecundity parameters used by PPA-SiBGC are easy to calculate using common field inventory data. PPA-SiBGC is simpler to transfer in this regard given the wide availability of forest inventory data.
3. *Size of the program* PPA-SiBGC is approximately 500 lines of R code, while LANDIS-II NECN is estimated at 0.5 million lines of C# code.
4. *Cross-platform support* While Linux support may soon be supported with Microsoft.NET Core, LANDIS-II NECN is written in C# and is thus limited to Microsoft Windows platforms. Meanwhile, PPA-SiBGC is written in standard R code and is fully cross-platform.
5. *Ease of training new users* While both models have a learning curve, the practical simplicity of PPA-SiBGC may make it easier to train new users. While LANDIS-II NECN contains more mechanistic processes and related parameters, these come at the cost of confusing new users. The model wrapper library we developed as part of this work vastly eases the operation of both models. Future studies should measure the time required for new users to effectively operate both models.

## 4. Discussion

The advancement of processor architectures has facilitated the development of increasingly complex forest models. Each new generation of processors allows researchers to conduct large-scale simulations faster and more efficiently than previous designs. As a result, forest models have grown into large, complex, analytically intractable programs. Rigorous intercomparison of models developed by different research groups, as well as the diagnosis of new versions of established models, is therefore a critical step in further advancing ecosystem models. This ensures that models are properly diagnosed and compared in a consistent, reliable, and transparent manner. Too often, model intercomparisons are conducted by each separate research group applying their own model in a manner that is, at best, inconsistent and opaque. In this work, we extended our model intercomparison by further providing wrapper functions that may be used to benchmark additional models or sites through a unified modeling framework. This ensures the consistency and transparency of intercomparison results.

The presented research is intended to establish the groundwork for future model intercomparison studies at both sites in order to advance the design of new models. Furthermore, we hope that this work will inspire a new generation of forest model intercomparisons in North America, which are sorely absent. Forest models have proven to be a critical testbed for improving the representation of vegetation dynamics in global terrestrial biosphere models [42,43,99], given the importance of forests in the global carbon cycle and the increased detail of local- to regional-scale models. Model benchmarking datasets and related results should be publicly shared and regularly updated with version-controlled software repositories (e.g., GitHub or GitLab), as is commonplace in the machine learning research community. Cloud computing providers may provide full reproducibility for cases where compute is limiting. In general, there is a broad disparity between modern software tools and existing forest models.

One important new forest model in development is a next-generation model from the SORTIE-PPA family of models, known as SORTIE-NG. This new model combines mechanistic representations of demographic processes, energetic and biogeochemical fluxes, and landscape disturbance dynamics, using hierarchical multiscale modeling with a modular component-based software framework [100]. Along with LM3-PPA [47], SORTIE-NG is among the first of a new class of hybrid models that we term ‘cohort-leaf’ models for their partitioning of energetic and biogeochemical fluxes amongst dynamic vegetation cohorts, instead of a single vertical ‘big-leaf’ profile. The SORTIE-NG model includes evolutionary optimality principles as well as phenotype plasticity and intraspecific genetic diversity through first-class support for probabilistic modeling, borrowing design principles from probabilistic programming languages (e.g., [101]). Thus, SORTIE-NG is intended to be the first forest model to bridge the divide between big-leaf, gap, and landscape models, and to be designed from the outset as a probabilistic modeling framework [100]. Future model extensions are in the planning stages, including the first machine-learning modeling interface included in an ecosystem model.

While implemented in a ‘close-to-metal’ language (i.e., C++17) and designed for efficiency, SORTIE-NG is vastly more computationally demanding than the PPA-SiBGC model used in this paper. Yet, we anticipate that SORTIE-NG will be able to improved the fidelity to observed fluxes, which is the major shortcoming of both models considered in this paper. Similarly, there is a new version of the LANDIS-II NECN model in development known as NECN-Hydro, which remains a simplified variant of the Century model, but includes more detailed hydrological processes. The currently presented work provides not only an intercomparison of two current state-of-art models, but also open-source software and wrapper functions for simple and rapid comparison of our results with new models or sites. The selected forested ecosystems modeled in this work are among the best-studied model forests on Earth today. Specifically, the EMS EC flux tower at Harvard Forest is the longest running flux tower in the United States. Extensions of the presented work will allow rigorous model comparison methodologies for forest models that will benefit the research community at large.

Extensions of this work may also address the robustness of model predictions to variations in parameter values. The parameterization of complex forest biogeochemistry models such as LANDIS-II NECN and PPA-SiBGC is an important problem for consideration. Models such as LANDIS-II NECN operate with an order of magnitude more parameters than PPA-SiBGC, which can each be estimated with different levels of accuracy. Often, we know only the range of parameter values while parameterization can also depend on the statistical approach employed. Meanwhile, authors routinely employ additional model calibration that consists of adjusting parameters in order to obtain improved fitness, which we explicitly avoided in this study.

Conducting such analyses through a unified software framework in a fully transparent and reproducible manner is therefore of the utmost importance. This is exactly the type of analyses that our provided software is designed to support. In a parallel line of research, we extend this base-level implementation into a generic application programming interface (API) and toolkit for geoscientific simulation models, known as Erde [102], supporting both R and Python. The Erde framework provides machine learning model emulation, robust loss estimation, parameter optimization, probabilistic parameterization, samplers such as Latin hypercube sampling and Markov Chain Monte Carlo, and a number of other helper methods designed for complex simulation models. We utilize the Erde framework in the design of Erde Gym, a toolkit for developing and comparing optimization algorithms in the geosciences with a focus on reinforcement learning [102]. For the first time, Erde Gym will allow us to model systems (e.g., evolutionary plant optimality) as intelligent agents able to navigate complex environments.

### 4.1. Limitations

This study, similar to most other modeling studies, was limited by the availability, quality, and quantity of observational data. The lack of temporal depth in this data poses substantial challenges in modeling the long-term effects of forest succession, as these processes can operate on a century timescale or longer. However, diagnosing succession was not the aim of this study, as we instead focus on near-term validation of forest models using field measurements and EC flux tower data. Another limitation is that these methods may be challenging to implement for sites that are less well-characterized, particularly in the absence of EC flux tower data and/or tree species parameters. A combination of tower-based and remote sensing observations may help overcome this challenge in the coming years with advances in machine learning. In addition, the poor performance of both tested models in capturing fluxes and excellent performance in capturing stocks indicate that the two current models should be applied in cases where stocks, rather than fluxes, are of primary interest.

### 4.2. Future opportunities

Future studies should expand upon the PPA with a first-principles representation of energetic and biogeochemical above- and below-ground processes in a modern component-based software framework. This work should fuse the new state-of-the-art forest biogeochemistry model with a model wrapper API written in R or Python, in order to expand native model functions to include Monte Carlo methods, machine-learning model emulation, robust loss functions, and optimization through a simple API enabling reproducibility. This would combine a high-performance forest model written in a compiled language with a simple, user-friendly interface written in an interpreted language, combining the best of both worlds. We are currently conducting work along this line by fusing the SORTIE-NG model with the Erde framework in order to develop state-of-the-art and user-friendly modeling capabilities, inspired by the design of modern deep learning frameworks such as PyTorch [103] and the Keras API [104].

In addition, there is a clear opportunity to link individual-based models such as PPA-SiBGC and SORTIE-NG to remote sensing data including airborne laser scanning or high-resolution multiview-stereo imagery (i.e., structure-from-motion), and hyperspectral indices of vegetation growth or stress. This line of work may assess opportunities for Bayesian data assimilation in addition to model parameterization and validation using detailed wall-to-wall forest structure maps. As models such as LES [30] provide more structural detail, spatially explicit data will be needed to parameterize the next generation of models. New data collection methods (e.g., [105]) will also be needed as the geometric realism of models advances toward the photorealistic detail offered by procedural models such as Lindenmayer- or L-systems [106,107].

## 5. Conclusions

In conclusion, the PPA-SiBGC and LANDIS-II NECN models represent vegetation dynamics previously absent in modeling studies at these sites. These include, “…long-term increases in tree biomass, successional change in forest composition, and disturbance events, processes not well represented in current models,” which drive interannual variation in NEE [55]. While the timescale of our simulations were decidedly short-term due to data limitations, both models showed good performance. While PPA-SiBGC showed stronger performance across the range of metrics tested, including the logistics of model deployment, LANDIS-II NECN also performed well across the metrics tested. Further studies are needed to compare more aspects of these and other models based on an array of performance criteria.

Ultimately, we hope that this study serves as the foundation for future forest ecosystem model intercomparisons for the North American continent, similar in spirit to the former TDE Ecosystem Model Intercomparison project [39]. This may help create the impetus for a Global Forest Model Intercomparison Project (ForestMIP) together with modeling groups on other continents. The aims of this research were not to determine which model is ‘best’ for prognosis at two locations, but to improve the capabilities of existing models across a range of locations in order to advance earth system models. In this regard, there are beneficial aspects to both modeling approaches and the trade-offs presented largely depend on the desired application. Counter to the classical modeling trade-off of Levins [36], improvements in precision and generality resulted from realism.

## Supplementary Materials

Parameter tables for both models and sites are provided in Appendix 5. All model, parameter, script files used in this model intercomparison exercise are available for download at the following public GitHub repository:

> https://github.com/adam-erickson/ecosystem-model-comparison

The repository provides tables containing parameter values and climate drivers used in the PPA-SiBGC and LANDIS-II NECN model simulations for the two model intercomparison sites. Tree species codes are adopted from the USDA PLANTS database, accessible at the following URL:

> https://plants.usda.gov

Scripts provided include a simple object-oriented forest biogeochemistry model wrapper library implemented in the R language [95]. The model wrapper library includes a number of features for simplifying the operation of this class of models, including functions for cleaning up and parsing model outputs into memory in a common format for comparison. Importantly, the wrapper library enables full reproducibility of results through the *hf_ems.r* and *jerc_rd.r* scripts. Using these scripts with the object-oriented *classes.r* model wrapper, it is possible to load pre-computed model results and calculate all intercomparison metrics for verification. The directory structure of the repository is shown in Figure 7.

**Figure 7.**
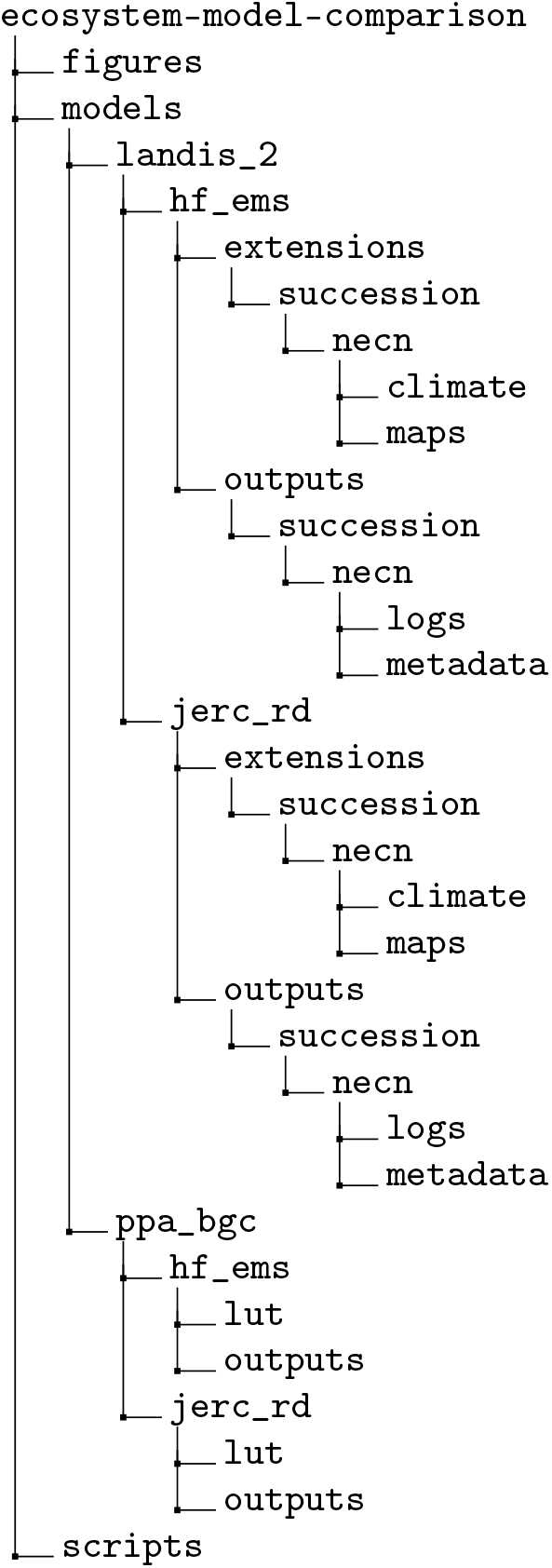
Directory structure of the GitHub repository

## Author Contributions

Individual contributions provided to complete this work include the following: conceptualization, N.S.; methodology, A.E. and N.S.; software, A.E.; validation, A.E.; formal analysis, A.E.; investigation, A.E.; resources, N.S.; data curation, A.E.; writing—original draft preparation, A.E.; writing—review and editing, A.E. and N.S.; visualization, A.E.; supervision, N.S.; project administration, N.S.; funding acquisition, N.S.

## Acknowledgments

We would like to thank U.S. Department of Defense, Army Corps of Engineers, and the Environmental Security Technology Certification Program (ESTCP) for providing support necessary to conduct this work. We also thank Harvard University and Jones Ecological Research Center for kindly providing data to conduct the analyses. We would further like to thank those that provided guidance on parameterization of the LANDIS-II model. We thank Drs. Louise Loudermilk of USDA Forest Service and Steven Flanagan of Tall Timbers Research Station for providing guidance on parameterization for the Red Dirt flux tower site at JERC, and Dr. Matthew Duveneck of New England Conservatory for providing guidance on parameterization for the HF-EMS EC flux tower site at Harvard Forest. We also acknowledge Drs. Melissa Lucash and Robert Scheller, who provided comments. Last, we would like to thank Drs. Bradley Case, Hannah Buckley, Audrey Barket-Plotkin, David Orwig, Aaron Ellison and Zachary Robbins for providing some of the crown allometry parameters for Harvard Forest.

## Abbreviations

The following abbreviations are used in this manuscript

ANPP: Aboveground net primary production
API: Application programming interface
BGC: Biogeochemistry
COST: Cooperation in Science and Technology
CPU: Central processing unit
CSV: Comma-separated values
DoD: Department of Defense
EC: Eddy covariance
ED: Ecosystem Demography model
EMS: Environmental Measurement Station
FVS: Forest Vegetation Simulator
GPGPU: General-purpose graphics processing unit
HF: Harvard Forest
IBIS2: Integrated Biosphere Simulator 2
JERC: Jones Ecological Research Center
L-systems: Lindenmayer systems
LANDIS-II: Landscape Disturbance and Succession model 2
LM3: Land Model 3
LPJ-GUESS: Lund-Potsdam-Jena General Ecosystem Simulator
MAE: Mean absolute error
MC1: MAPSS-Century-1 model
NECN: Net Ecosystem Carbon and Nitrogen model
NEE: Net ecosystem exchange
NSE: Nash-Sutcliffe efficiency
PPA: Perfect Plasticity Approximation model
ProFoUnd: Towards robust projections of European forests under climate change
RAM: Random access memory
RD: Red Dirt
RMSE: Root mean squared error
SAS: Size- and age-structured equations
SOC: Soil organic carbon
SON: Soil organic nitrogen
TDE: Throughfall Displacement Experiment

## Appendix A. Eddy covariance flux tower measurements

### Appendix A.1. HF-EMS EC Flux Tower

Recent historical mean daily fluxes of temperature (° *C*), ecosystem respiration (*μmol CO*_2_ *m*^−2^), and NEE (*μmol C m*^−2^) for the HF-EMS tower are shown in Figure A1.

**Figure A1.**
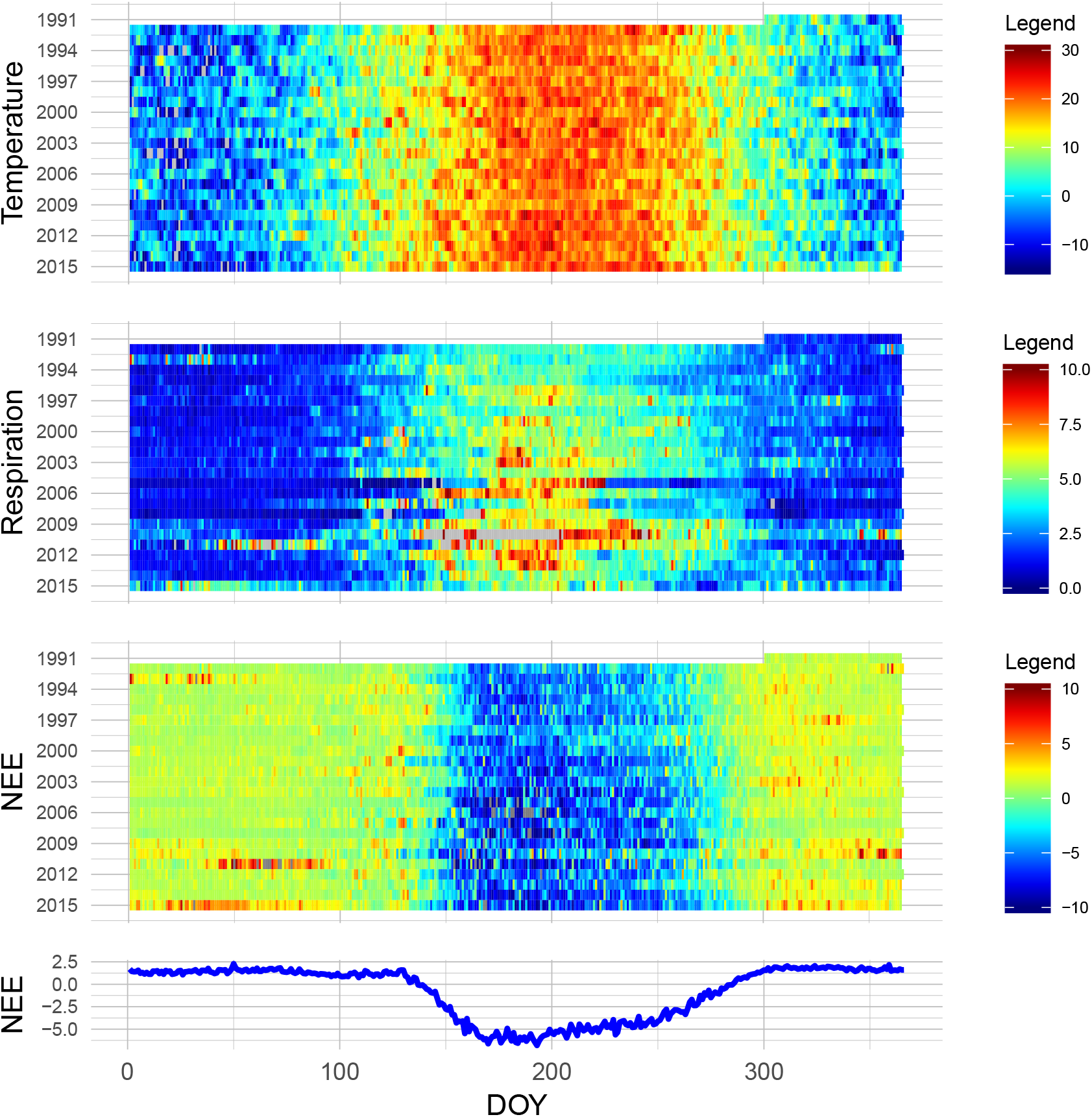
HF-EMS tower daily averages

Patterns in daytime and nighttime NEE are shown in Figure A2. This was calculated by taking daily mean NEE values for three-hour windows surrounding noon and midnight, respectively (1100-1300 and 2300-0100 hours). These patterns are important to diagnose, as they demonstrate responses to a gradient of light and temperature conditions.

**Figure A2.**
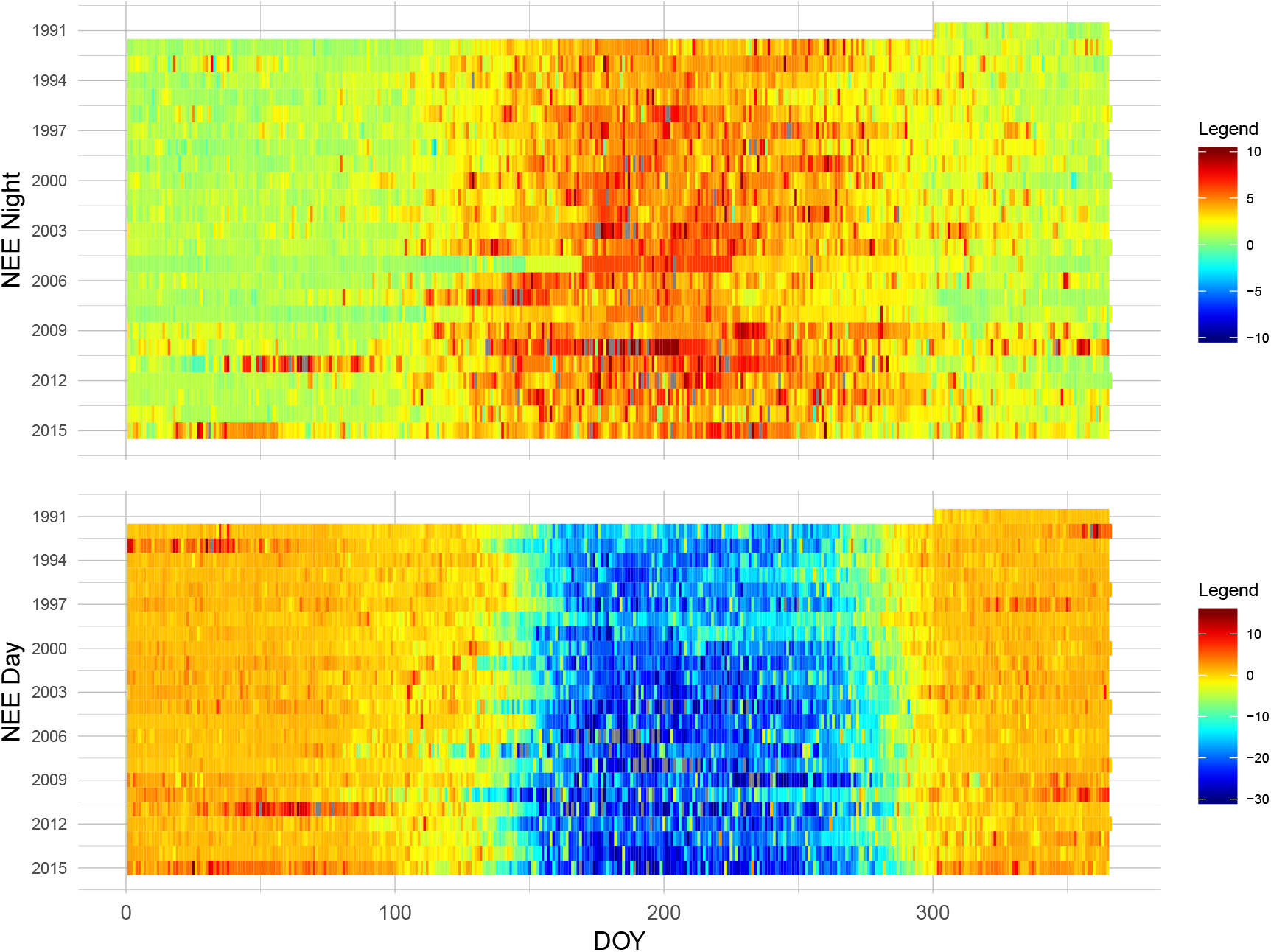
HF-EMS tower daily diurnal averages

### Appendix A.2. JERC-RD EC Flux Tower

Recent historical mean daily fluxes of latent heat flux (LE) (*W m*^−2^), ecosystem respiration (*μmol CO*_2_ *m*^−2^), and NEE (*μmol C m*^−2^) for the RD flux tower are shown in Figure A3.

**Figure A3.**
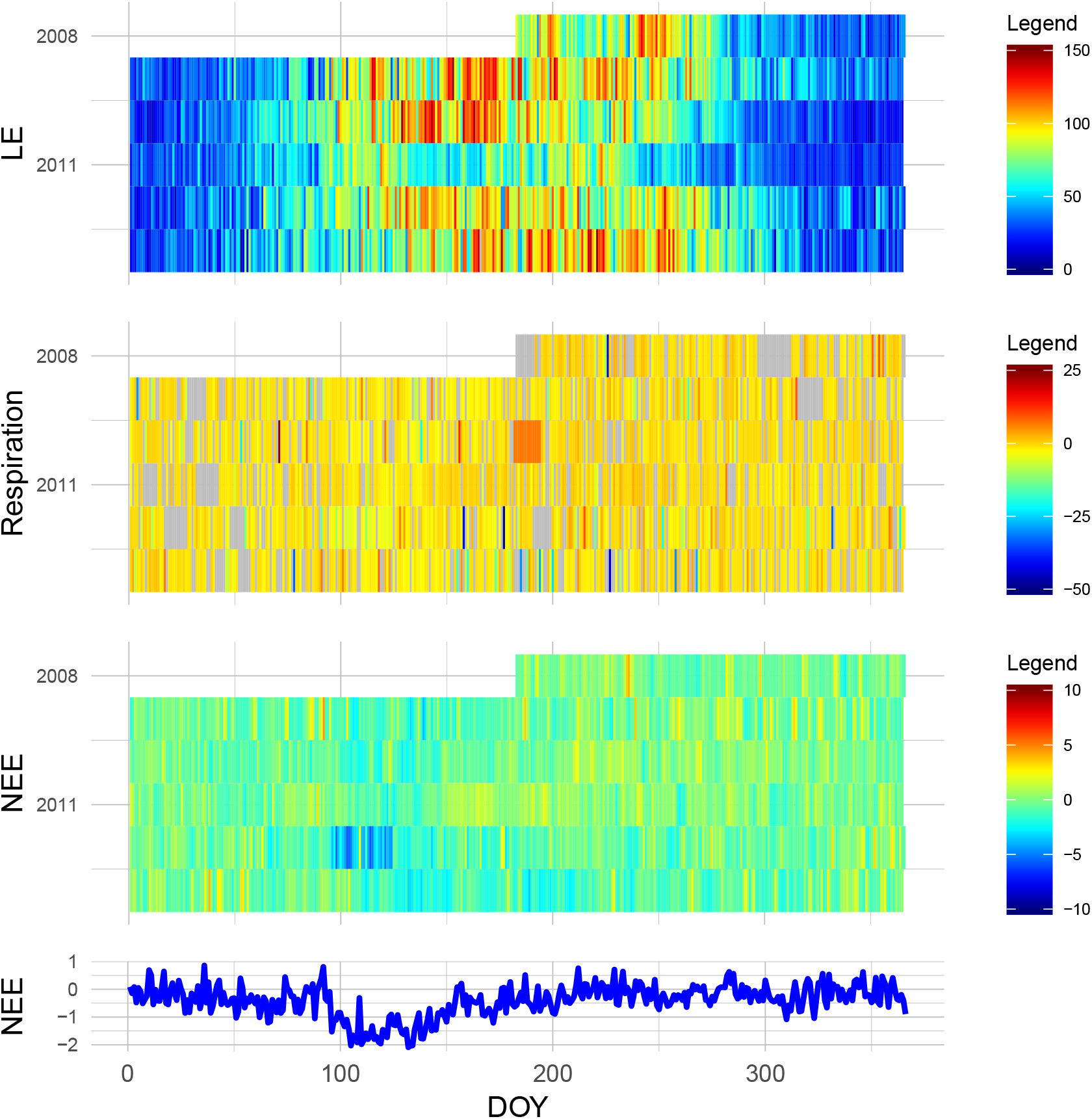
JERC-RD tower daily averages

Patterns of daytime and nighttime NEE are shown in Figure A4. Again, this was calculated by taking daily mean NEE values for three-hour windows surrounding noon and midnight, respectively (1100-1300 and 2300-0100 hours).

**Figure A4.**
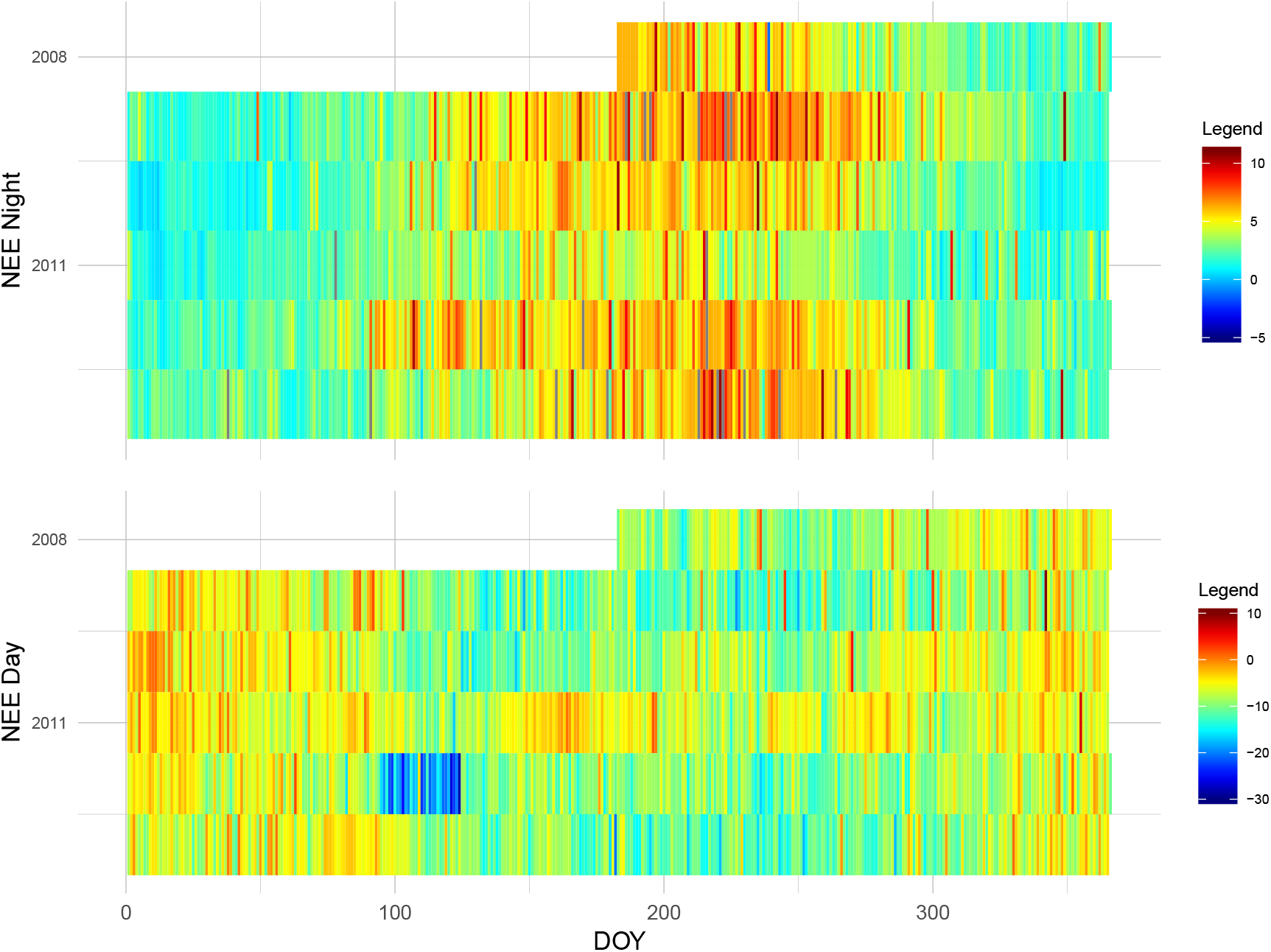
JERC-RD tower daily diurnal averages

## Appendix B. Site maps

Below, we provide maps of the two research sites for reference. First is the HF-EMS EC flux tower with landcover classes Figure A5.

**Figure A5.**
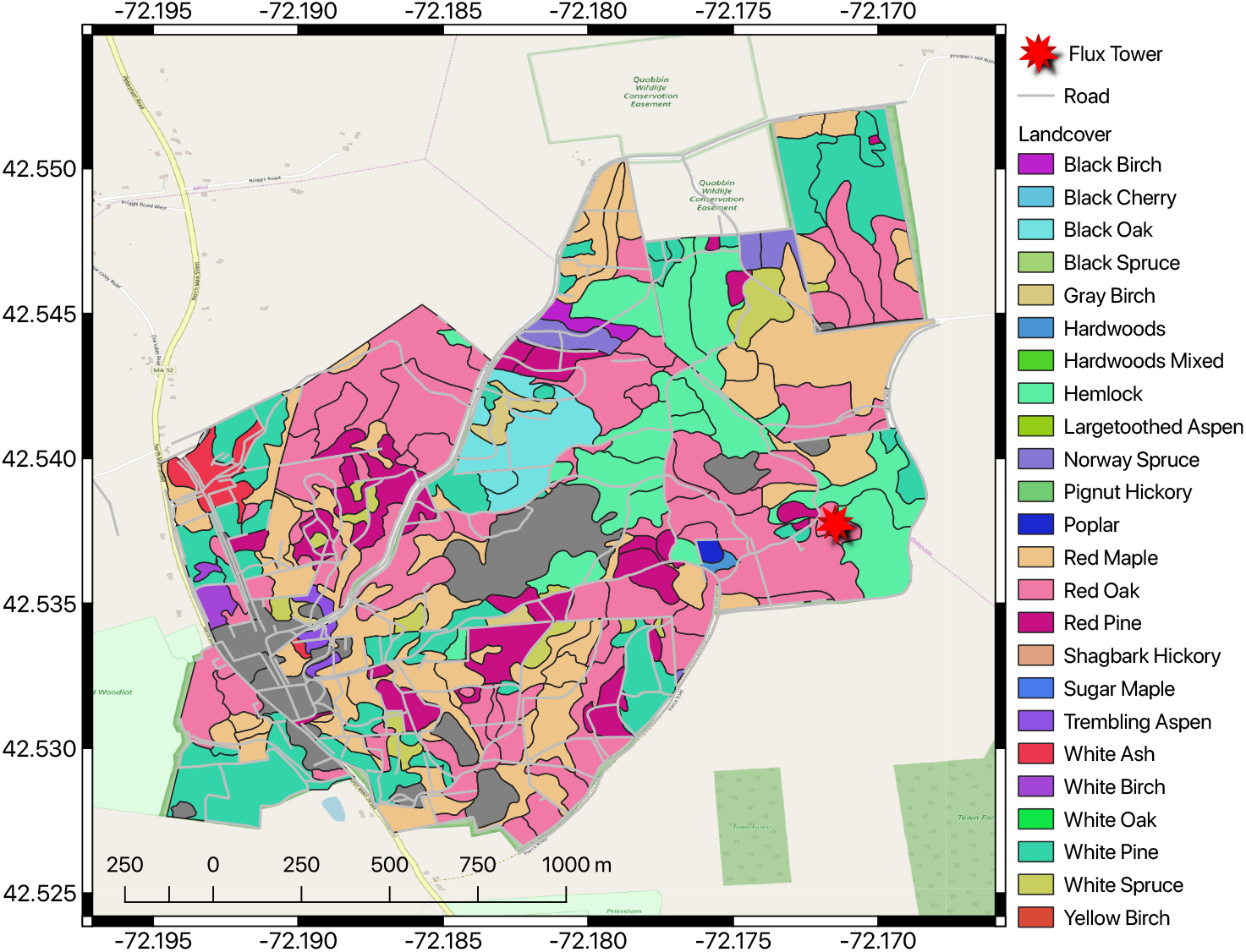
HF-EMS flux tower and landcover classes

Next is the JERC-RD flux tower with landcover classes Figure A6.

**Figure A6.**
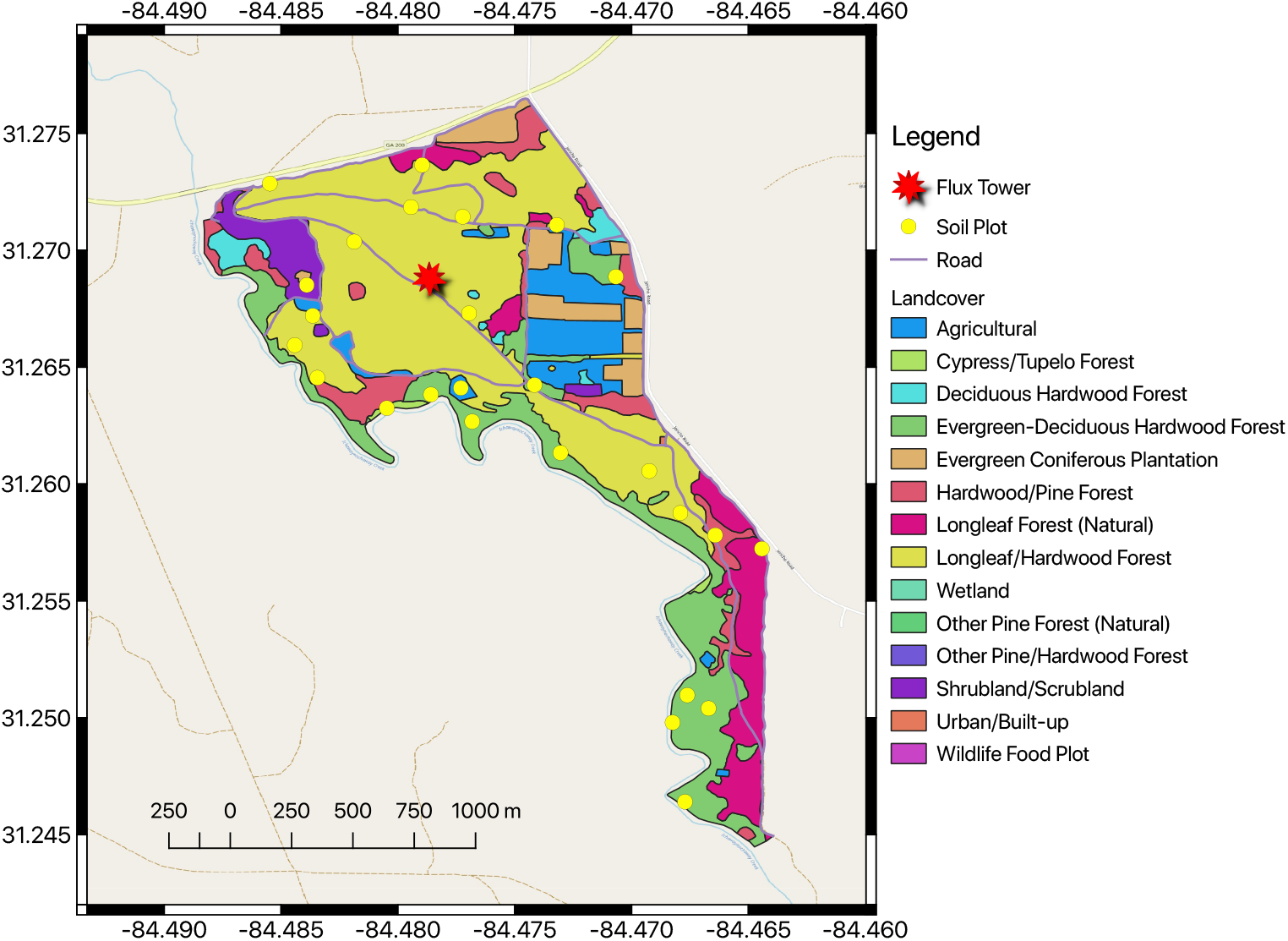
JERC-RD flux tower and landcover classes

## Appendix C. Model Parameters

### Appendix C.1. HF-EMS

#### Appendix C.1.1. PPA-SiBGC

**Table A1.**
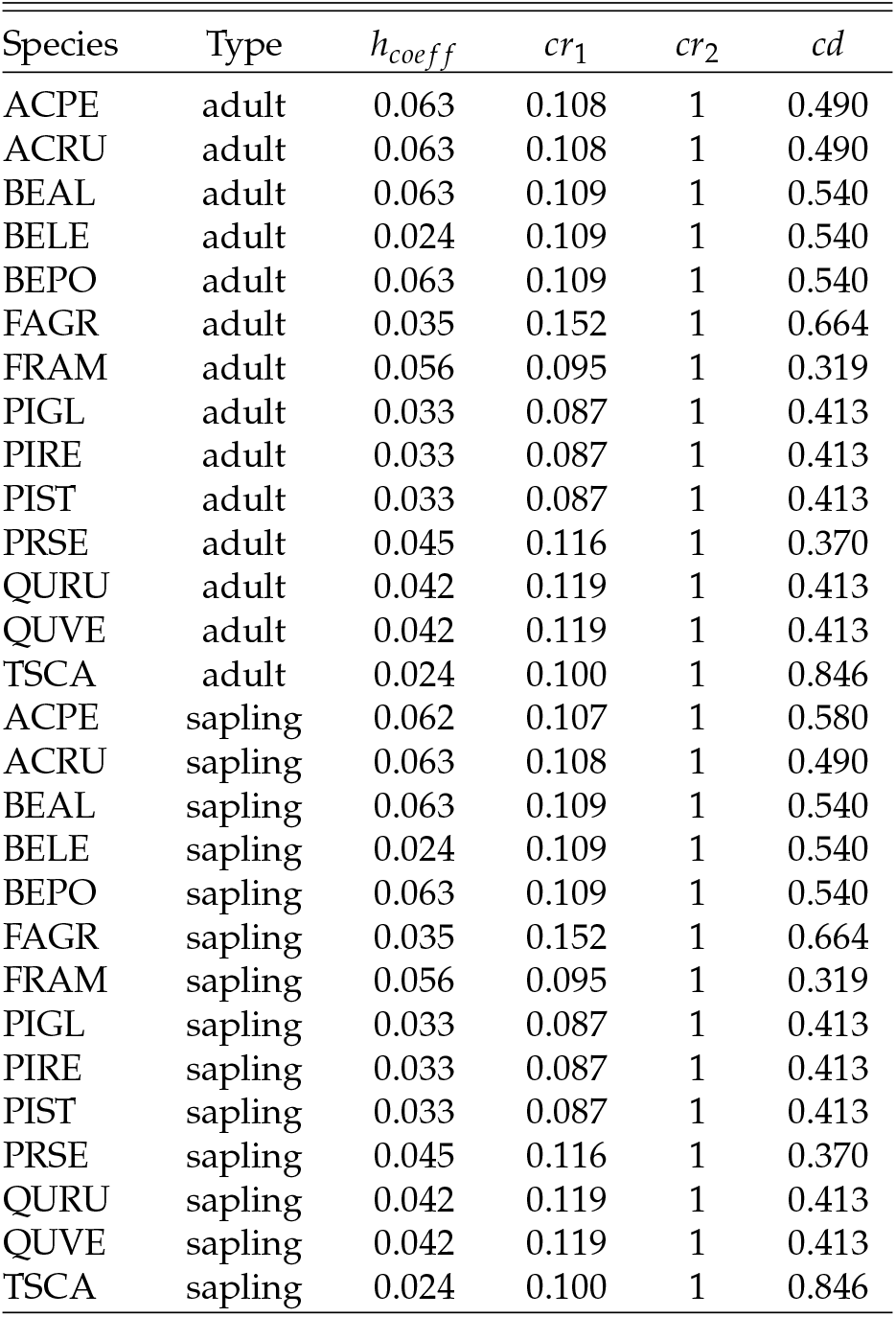
Species crown allometry parameters

| Species | Type | $h_{coeff}$ | $cr_1$ | $cr_2$ | $cd$ |
| --- | --- | --- | --- | --- | --- |
| ACPE | adult | 0.063 | 0.108 | 1 | 0.490 |
| ACRU | adult | 0.063 | 0.108 | 1 | 0.490 |
| BEAL | adult | 0.063 | 0.109 | 1 | 0.540 |
| BELE | adult | 0.024 | 0.109 | 1 | 0.540 |
| BEPO | adult | 0.063 | 0.109 | 1 | 0.540 |
| FAGR | adult | 0.035 | 0.152 | 1 | 0.664 |
| FRAM | adult | 0.056 | 0.095 | 1 | 0.319 |
| PIGL | adult | 0.033 | 0.087 | 1 | 0.413 |
| PIRE | adult | 0.033 | 0.087 | 1 | 0.413 |
| PIST | adult | 0.033 | 0.087 | 1 | 0.413 |
| PRSE | adult | 0.045 | 0.116 | 1 | 0.370 |
| QURU | adult | 0.042 | 0.119 | 1 | 0.413 |
| QUVE | adult | 0.042 | 0.119 | 1 | 0.413 |
| TSCA | adult | 0.024 | 0.100 | 1 | 0.846 |
| ACPE | sapling | 0.062 | 0.107 | 1 | 0.580 |
| ACRU | sapling | 0.063 | 0.108 | 1 | 0.490 |
| BEAL | sapling | 0.063 | 0.109 | 1 | 0.540 |
| BELE | sapling | 0.024 | 0.109 | 1 | 0.540 |
| BEPO | sapling | 0.063 | 0.109 | 1 | 0.540 |
| FAGR | sapling | 0.035 | 0.152 | 1 | 0.664 |
| FRAM | sapling | 0.056 | 0.095 | 1 | 0.319 |
| PIGL | sapling | 0.033 | 0.087 | 1 | 0.413 |
| PIRE | sapling | 0.033 | 0.087 | 1 | 0.413 |
| PIST | sapling | 0.033 | 0.087 | 1 | 0.413 |
| PRSE | sapling | 0.045 | 0.116 | 1 | 0.370 |
| QURU | sapling | 0.042 | 0.119 | 1 | 0.413 |
| QUVE | sapling | 0.042 | 0.119 | 1 | 0.413 |
| TSCA | sapling | 0.024 | 0.100 | 1 | 0.846 |

**Table A2.** Species biomass equation parameters

| Species | $b_0$ | $b_1$ | $f_{stem}$ | $f_{branch}$ | $f_{leaf}$ | $f_{root}$ | $f_{soil}$ |
| --- | --- | --- | --- | --- | --- | --- | --- |
| ACPE | −2.047 | 2.385 | 0.700 | 0.230 | 0.070 | 0.240 | 0.680 |
| ACRU | −2.047 | 2.385 | 0.700 | 0.230 | 0.070 | 0.240 | 0.680 |
| BEAL | −1.810 | 2.348 | 0.700 | 0.230 | 0.070 | 0.240 | 0.680 |
| BELE | −1.810 | 2.348 | 0.700 | 0.230 | 0.070 | 0.240 | 0.680 |
| BEPO | −2.227 | 2.451 | 0.700 | 0.230 | 0.070 | 0.240 | 0.680 |
| FAGR | −2.070 | 2.441 | 0.700 | 0.230 | 0.070 | 0.240 | 0.680 |
| FRAM | −1.838 | 2.352 | 0.700 | 0.230 | 0.070 | 0.240 | 0.680 |
| PIGL | −2.136 | 2.323 | 0.700 | 0.230 | 0.070 | 0.240 | 0.680 |
| PIRE | −2.618 | 2.464 | 0.700 | 0.230 | 0.070 | 0.240 | 0.680 |
| PIST | −2.618 | 2.464 | 0.700 | 0.230 | 0.070 | 0.240 | 0.680 |
| PRSE | −2.212 | 2.413 | 0.700 | 0.230 | 0.070 | 0.240 | 0.680 |
| QURU | −2.070 | 2.441 | 0.700 | 0.230 | 0.070 | 0.240 | 0.680 |
| QUVE | −2.070 | 2.441 | 0.700 | 0.230 | 0.070 | 0.240 | 0.680 |
| TSCA | −2.348 | 2.388 | 0.700 | 0.230 | 0.070 | 0.240 | 0.680 |

**Table A3.** Biomass carbon fraction parameters

| $f_{stem}$ | $f_{branch}$ | $f_{leaf}$ | $f_{root}$ | $f_{soil}$ |
| --- | --- | --- | --- | --- |
| 0.500 | 0.500 | 0.500 | 0.500 | 0.143 |

**Table A4.** Species DBH increment parameters

| Species | Type | $I_{DBH}$ |
| --- | --- | --- |
| ACPE | adult | 0.277 |
| ACRU | adult | 0.312 |
| BEAL | adult | 0.280 |
| BELE | adult | 0.198 |
| BEPO | adult | 0.103 |
| FAGR | adult | 0.303 |
| FRAM | adult | 0.149 |
| PIGL | adult | 0.274 |
| PIRE | adult | 0.390 |
| PIST | adult | 0.277 |
| PRSE | adult | 0.120 |
| QURU | adult | 0.420 |
| QUVE | adult | 0.322 |
| TSCA | adult | 0.563 |
| ACPE | sapling | 0.895 |
| ACRU | sapling | 0.269 |
| BEAL | sapling | 0.520 |
| BELE | sapling | 0.201 |
| BEPO | sapling | 0.300 |
| FAGR | sapling | 0.530 |
| FRAM | sapling | 0.500 |
| PIGL | sapling | 0.353 |
| PIRE | sapling | 0.350 |
| PIST | sapling | 0.350 |
| PRSE | sapling | 0.200 |
| QURU | sapling | 0.098 |
| QUVE | sapling | 0.100 |
| TSCA | sapling | 0.509 |

**Table A5.** Species mortality parameters

| Species | Type | $p_{mortality}$ |
| --- | --- | --- |
| ACPE | adult | 0.115 |
| ACRU | adult | 0.030 |
| BEAL | adult | 0.035 |
| BELE | adult | 0.009 |
| BEPO | adult | 0.032 |
| FAGR | adult | 0.015 |
| FRAM | adult | 0.004 |
| PIGL | adult | 0.074 |
| PIRE | adult | 0.023 |
| PIST | adult | 0.010 |
| PRSE | adult | 0.009 |
| QURU | adult | 0.007 |
| QUVE | adult | 0.001 |
| TSCA | adult | 0.022 |
| ACPE | sapling | 0.001 |
| ACRU | sapling | 0.873 |
| BEAL | sapling | 0.001 |
| BELE | sapling | 0.667 |
| BEPO | sapling | 0.001 |
| FAGR | sapling | 0.354 |
| FRAM | sapling | 0.001 |
| PIGL | sapling | 0.001 |
| PIRE | sapling | 0.001 |
| PIST | sapling | 0.001 |
| PRSE | sapling | 0.001 |
| QURU | sapling | 0.001 |
| QUVE | sapling | 0.001 |
| TSCA | sapling | 0.821 |

**Table A6.**
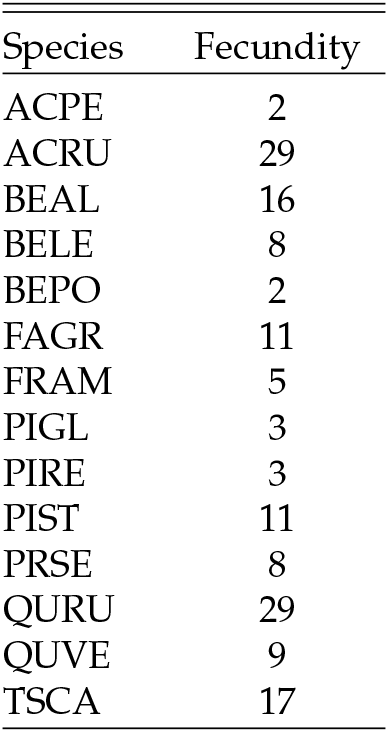
Species fecundity parameters

**Table A7.** Species C:N ratio parameters

| Species | $CN_{stem}$ | $CN_{branch}$ | $CN_{leaf}$ | $CN_{litter}$ | $CN_{root}$ | $CN_{soil}$ |
| --- | --- | --- | --- | --- | --- | --- |
| ACPE | 548.590 | 71.460 | 30.460 | 58.800 | 68.548 | 23.087 |
| ACRU | 548.590 | 71.460 | 30.460 | 58.800 | 68.548 | 23.087 |
| BEAL | 548.590 | 71.460 | 22.420 | 58.800 | 68.548 | 23.087 |
| BELE | 548.590 | 71.460 | 21.200 | 58.800 | 68.548 | 23.087 |
| BEPO | 548.590 | 71.460 | 21.560 | 58.800 | 68.548 | 23.087 |
| FAGR | 548.590 | 71.460 | 22.420 | 58.800 | 68.548 | 23.087 |
| FRAM | 548.590 | 71.460 | 21.910 | 58.800 | 68.548 | 23.087 |
| PIGL | 548.590 | 71.460 | 38 | 58.800 | 68.548 | 23.087 |
| PIRE | 548.590 | 71.460 | 33 | 58.800 | 68.548 | 23.087 |
| PIST | 548.590 | 71.460 | 38 | 58.800 | 68.548 | 23.087 |
| PRSE | 548.590 | 71.460 | 21.500 | 58.800 | 68.548 | 23.087 |
| QURU | 548.590 | 71.460 | 21.920 | 58.800 | 68.548 | 23.087 |
| QUVE | 548.590 | 71.460 | 21.920 | 58.800 | 68.548 | 23.087 |
| TSCA | 548.590 | 71.460 | 42.520 | 58.800 | 68.548 | 23.087 |

#### Appendix C.1.2. LANDIS-II NECN

**Table A8.** NECN adjustment parameters

| Parameter | Value |
| --- | --- |
| $p_{est}$ modifier | 0.1 |
| $N_{mineral}$ initial | 3.0 |
| $Fuels_{fine}$ initial | 0.1 |
| $N_{atmos}$ slope | 0.007 |
| $N_{atmos}$ intercept | 0.011 |
| $Latitude_{deg}$ | 43.3 |
| $r_{denitrification}$ | 0.001 |
| $r_{decay}$ surface | 0.65 |
| $r_{decay}$ SOM1 | 1.0 |
| $r_{decay}$ SOM2 | 0.125 |
| $r_{decay}$ SOM3 | 0.0002 |

**Table A9.** NECN maximum LAI parameters

| $Class_{shade}$ | $LAI_{max}$ |
| --- | --- |
| 1 | 1 |
| 2 | 2.5 |
| 3 | 3.5 |
| 4 | 6 |
| 5 | 8 |

**Table A10.** NECN light establishment parameters

| $Class_{shade}$ | $Shade_0$ | $Shade_1$ | $Shade_2$ | $Shade_3$ | $Shade_4$ | $Shade_5$ |
| --- | --- | --- | --- | --- | --- | --- |
| 1 | 1 | 1 | 0.25 | 0.1 | 0 | 0 |
| 2 | 0.5 | 0.5 | 1 | 0.25 | 0.1 | 0 |
| 3 | 0.1 | 0.5 | 1 | 1 | 0.5 | 0.1 |
| 4 | 0.1 | 0.25 | 0.5 | 0.5 | 1 | 0.25 |
| 5 | 0 | 0.1 | 0.25 | 0.25 | 0.5 | 1 |

**Table A11.**
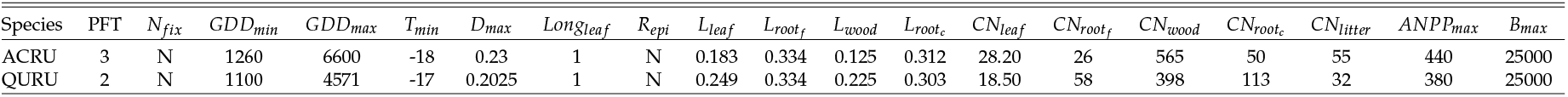
NECN species parameters

**Table A12.** Functional group parameters

| PFT | Index | $T_{mean}$ | $T_{max}$ | $T_{shape}$ | $T_{shape}$ | $f_{cf}$ | $BTOLAI$ | $kLAI$ | $LAI_{max}$ | $PPRPTS_2$ | $PPRPTS_3$ | $r_{decayw}$ | $m_{wood}$ | $m_{shape}$ | $drop_{month}$ | $f_{rootc}$ | $f_{rootf}$ |
| --- | --- | --- | --- | --- | --- | --- | --- | --- | --- | --- | --- | --- | --- | --- | --- | --- | --- |
| Oaks | 2 | 25 | 40 | 1.5 | 2.5 | 0.6 | -0.9 | 10000 | 9 | 0.1 | 0.8 | 0.5 | 0.0006 | 15 | 9 | 0.2 | 0.5 |
| NorthHardwoods | 3 | 25 | 40 | 1.5 | 2.5 | 0.6 | -0.9 | 7000 | 10 | 1.5 | 0.96 | 0.7 | 0.0006 | 15 | 9 | 0.2 | 0.5 |

**Table A13.** Fire reduction parameters; inactive

| $Class_{severity}$ | $Reduction_{wood}$ | $Reduction_{litter}$ | $Reduction_{SOM}$ |
| --- | --- | --- | --- |
| 1 | 0.0 | 0.5 | 1.0 |
| 2 | 0.05 | 0.75 | 1.0 |
| 3 | 0.2 | 1.0 | 1.0 |
| 4 | 0.5 | 1.0 | 1.0 |
| 5 | 0.8 | 1.0 | 1.0 |

**Table A14.** Harvest reduction parameters; inactive

| Class | $Reduction_{wood}$ | $Reduction_{litter}$ | $Reduction_{SOM}$ | $Removal_{leaf}$ | $Removal_{wood}$ |
| --- | --- | --- | --- | --- | --- |
| HandThinning | 0.05 | 1.0 | 1.0 | 1.0 | 1.0 |
| MechThinning | 0.05 | 1.0 | 1.0 | 0.85 | 1.0 |

**Table A15.**
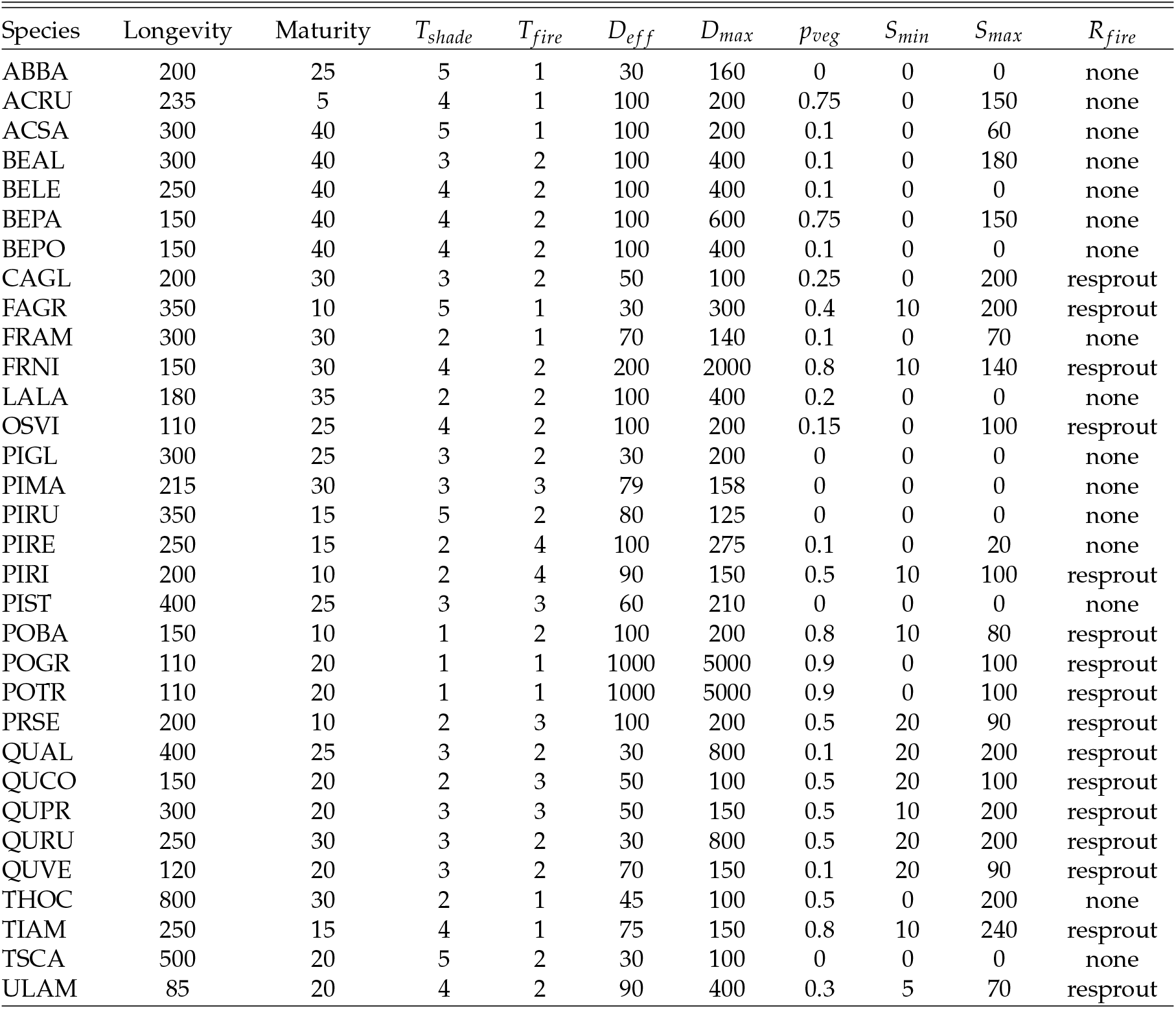
Species parameters; only ACRU and QURU were simulated

| Species | Longevity | Maturity | $T_{shade}$ | $T_{fire}$ | $D_{eff}$ | $D_{max}$ | $p_{veg}$ | $S_{min}$ | $S_{max}$ | $R_{fire}$ |
| --- | --- | --- | --- | --- | --- | --- | --- | --- | --- | --- |
| ABBA | 200 | 25 | 5 | 1 | 30 | 160 | 0 | 0 | 0 | none |
| ACRU | 235 | 5 | 4 | 1 | 100 | 200 | 0.75 | 0 | 150 | none |
| ACSA | 300 | 40 | 5 | 1 | 100 | 200 | 0.1 | 0 | 60 | none |
| BEAL | 300 | 40 | 3 | 2 | 100 | 400 | 0.1 | 0 | 180 | none |
| BELE | 250 | 40 | 4 | 2 | 100 | 400 | 0.1 | 0 | 0 | none |
| BEPA | 150 | 40 | 4 | 2 | 100 | 600 | 0.75 | 0 | 150 | none |
| BEPO | 150 | 40 | 4 | 2 | 100 | 400 | 0.1 | 0 | 0 | none |
| CAGL | 200 | 30 | 3 | 2 | 50 | 100 | 0.25 | 0 | 200 | resprout |
| FAGR | 350 | 10 | 5 | 1 | 30 | 300 | 0.4 | 10 | 200 | resprout |
| FRAM | 300 | 30 | 2 | 1 | 70 | 140 | 0.1 | 0 | 70 | none |
| FRNI | 150 | 30 | 4 | 2 | 200 | 2000 | 0.8 | 10 | 140 | resprout |
| LALA | 180 | 35 | 2 | 2 | 100 | 400 | 0.2 | 0 | 0 | none |
| OSVI | 110 | 25 | 4 | 2 | 100 | 200 | 0.15 | 0 | 100 | resprout |
| PIGL | 300 | 25 | 3 | 2 | 30 | 200 | 0 | 0 | 0 | none |
| PIMA | 215 | 30 | 3 | 3 | 79 | 158 | 0 | 0 | 0 | none |
| PIRU | 350 | 15 | 5 | 2 | 80 | 125 | 0 | 0 | 0 | none |
| PIRE | 250 | 15 | 2 | 4 | 100 | 275 | 0.1 | 0 | 20 | none |
| PIRI | 200 | 10 | 2 | 4 | 90 | 150 | 0.5 | 10 | 100 | resprout |
| PIST | 400 | 25 | 3 | 3 | 60 | 210 | 0 | 0 | 0 | none |
| POBA | 150 | 10 | 1 | 2 | 100 | 200 | 0.8 | 10 | 80 | resprout |
| POGR | 110 | 20 | 1 | 1 | 1000 | 5000 | 0.9 | 0 | 100 | resprout |
| POTR | 110 | 20 | 1 | 1 | 1000 | 5000 | 0.9 | 0 | 100 | resprout |
| PRSE | 200 | 10 | 2 | 3 | 100 | 200 | 0.5 | 20 | 90 | resprout |
| QUAL | 400 | 25 | 3 | 2 | 30 | 800 | 0.1 | 20 | 200 | resprout |
| QUCO | 150 | 20 | 2 | 3 | 50 | 100 | 0.5 | 20 | 100 | resprout |
| QUPR | 300 | 20 | 3 | 3 | 50 | 150 | 0.5 | 10 | 200 | resprout |
| QURU | 250 | 30 | 3 | 2 | 30 | 800 | 0.5 | 20 | 200 | resprout |
| QUVE | 120 | 20 | 3 | 2 | 70 | 150 | 0.1 | 20 | 90 | resprout |
| THOC | 800 | 30 | 2 | 1 | 45 | 100 | 0.5 | 0 | 200 | none |
| TIAM | 250 | 15 | 4 | 1 | 75 | 150 | 0.8 | 10 | 240 | resprout |
| TSCA | 500 | 20 | 5 | 2 | 30 | 100 | 0 | 0 | 0 | none |
| ULAM | 85 | 20 | 4 | 2 | 90 | 400 | 0.3 | 5 | 70 | resprout |

### Appendix C.2. JERC-RD

#### Appendix C.2.1. PPA-SiBGC

**Table A16.** Species crown allometry parameters

| Species | Type | $h_{coeff}$ | $cr1$ | $cr2$ | $cd$ |
| --- | --- | --- | --- | --- | --- |
| PIPA | adult | 0.033 | 0.087 | 1 | 0.413 |
| QUIN | adult | 0.042 | 0.119 | 1 | 0.413 |
| QUNI | adult | 0.042 | 0.119 | 1 | 0.413 |
| QUVI | adult | 0.042 | 0.119 | 1 | 0.413 |
| PIPA | sapling | 0.033 | 0.087 | 1 | 0.413 |
| QUIN | sapling | 0.042 | 0.119 | 1 | 0.413 |
| QUNI | sapling | 0.042 | 0.119 | 1 | 0.413 |
| QUVI | sapling | 0.042 | 0.119 | 1 | 0.413 |

**Table A17.** Species biomass equation parameters

| Species | $b_0$ | $b_1$ | $f_{stem}$ | $f_{branch}$ | $f_{leaf}$ | $f_{root}$ | $f_{soil}$ |
| --- | --- | --- | --- | --- | --- | --- | --- |
| PIPA | -3.051 | 2.647 | 0.700 | 0.230 | 0.070 | 0.240 | 0.680 |
| QUIN | -2.070 | 2.441 | 0.700 | 0.230 | 0.070 | 0.240 | 0.680 |
| QUNI | -2.070 | 2.441 | 0.700 | 0.230 | 0.070 | 0.240 | 0.680 |
| QUVI | -2.070 | 2.441 | 0.700 | 0.230 | 0.070 | 0.240 | 0.680 |

**Table A18.** Biomass carbon fraction parameters

| $f_{stem}$ | $f_{branch}$ | $f_{leaf}$ | $f_{root}$ | $f_{soil}$ |
| --- | --- | --- | --- | --- |
| 0.500 | 0.500 | 0.500 | 0.500 | 0.143 |

**Table A19.** Species DBH increment parameters

| Species | Type | $I_{DBH}$ |
| --- | --- | --- |
| PIPA | adult | 0.261 |
| QUIN | adult | 0.119 |
| QUNI | adult | 0.994 |
| QUVI | adult | 0.276 |
| PIPA | sapling | 0.197 |
| QUIN | sapling | 0.100 |
| QUNI | sapling | 0.440 |
| QUVI | sapling | 0.271 |

**Table A20.** Species mortality parameters

| Species | Type | $p_{mortality}$ |
| --- | --- | --- |
| PIPA | adult | 0.001 |
| QUIN | adult | 0.001 |
| QUNI | adult | 0.001 |
| QUVI | adult | 0.001 |
| PIPA | sapling | 0.174 |
| QUIN | sapling | 0.333 |
| QUNI | sapling | 0.143 |
| QUVI | sapling | 0.111 |

**Table A21.**
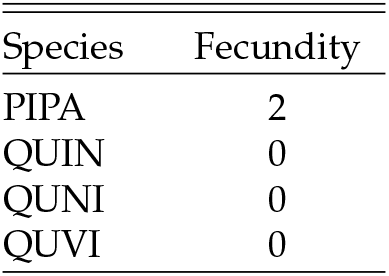
Species fecundity parameters

**Table A22.**
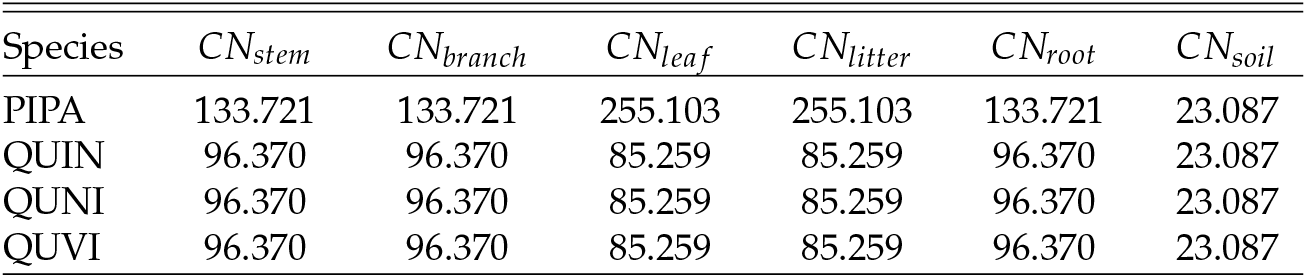
Species C:N ratio parameters

| Species | $CN_{stem}$ | $CN_{branch}$ | $CN_{leaf}$ | $CN_{litter}$ | $CN_{root}$ | $CN_{soil}$ |
| --- | --- | --- | --- | --- | --- | --- |
| PIPA | 133.721 | 133.721 | 255.103 | 255.103 | 133.721 | 23.087 |
| QUIN | 96.370 | 96.370 | 85.259 | 85.259 | 96.370 | 23.087 |
| QUNI | 96.370 | 96.370 | 85.259 | 85.259 | 96.370 | 23.087 |
| QUVI | 96.370 | 96.370 | 85.259 | 85.259 | 96.370 | 23.087 |

#### Appendix C.2.2. LANDIS-II NECN

**Table A23.** NECN adjustment parameters

| Parameter | Value |
| --- | --- |
| $p_{est}$ modifier | 0.4 |
| $N_{mineral}$ initial | 0.5 |
| $Fuels_{fine}$ initial | 0.1 |
| $N_{atmos}$ slope | 0.004 |
| $N_{atmos}$ intercept | 0.017 |
| $Latitude_{deg}$ | 31.220731 |
| $r_{denitrification}$ | 0.02 |
| $r_{decay}$ surface | 0.70 |
| $r_{decay}$ SOM1 | 0.81 |
| $r_{decay}$ SOM2 | 0.05 |
| $r_{decay}$ SOM3 | 0.00006 |

**Table A24.**
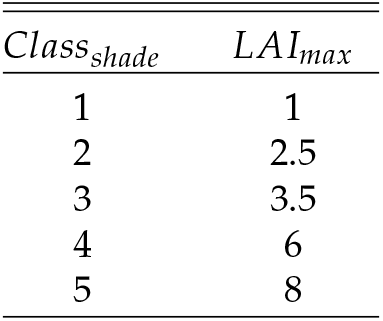
NECN maximum LAI parameters

**Table A25.** NECN light establishment parameters

| $Class_{shade}$ | $Shade_0$ | $Shade_1$ | $Shade_2$ | $Shade_3$ | $Shade_4$ | $Shade_5$ |
| --- | --- | --- | --- | --- | --- | --- |
| 1 | 1 | 1 | 0.25 | 0.1 | 0 | 0 |
| 2 | 0.5 | 0.5 | 1 | 0.25 | 0.1 | 0 |
| 3 | 0.1 | 1 | 1 | 1 | 0.5 | 0.1 |
| 4 | 0.1 | 0.25 | 0.5 | 0.5 | 1 | 0.25 |
| 5 | 0 | 0.1 | 0.25 | 0.25 | 0.5 | 1 |

**Table A26.** NECN species parameters

| Species | PFT | $N_{fix}$ | $GDD_{min}$ | $GDD_{max}$ | $T_{min}$ | $D_{max}$ | $Longleaf$ | $R_{epi}$ | $L_{leaf}$ | $L_{root_f}$ | $L_{wood}$ | $L_{root_c}$ | $CN_{leaf}$ | $CN_{root_f}$ | $CN_{wood}$ | $CN_{root_c}$ | $CN_{litter}$ | $ANPP_{max}$ | $B_{max}$ |
| --- | --- | --- | --- | --- | --- | --- | --- | --- | --- | --- | --- | --- | --- | --- | --- | --- | --- | --- | --- |
| QUIN | 2 | N | 3915 | 7000 | 1 | 0.423 | 1 | N | 0.293 | 0.23 | 0.23 | 0.35 | 24 | 48 | 500 | 333 | 55 | 250 | 15000 |
| QULA | 2 | N | 3915 | 7000 | 1 | 0.423 | 1 | N | 0.293 | 0.23 | 0.23 | 0.35 | 24 | 48 | 500 | 333 | 55 | 250 | 15000 |
| PIPA | 1 | N | 3915 | 7000 | 1 | 0.423 | 2 | N | 0.2 | 0.2 | 0.35 | 0.35 | 50 | 50 | 380 | 170 | 100 | 500 | 15000 |

**Table A27.** Functional group parameters

| PFT | Index | $T_{mean}$ | $T_{max}$ | $T_{shape}$ | $T_{shape}$ | $f_{C_f}$ | $BTOLAI$ | $kLAI$ | $LAI_{max}$ | $PPRPTS_2$ | $PPRPTS_3$ | $r_{decay_w}$ | $m_{wood}$ | $m_{shape}$ | $month_{drop}$ | $f_{root_c}$ | $f_{root_f}$ |
| --- | --- | --- | --- | --- | --- | --- | --- | --- | --- | --- | --- | --- | --- | --- | --- | --- | --- |
| Pine | 1 | 28 | 45 | 3.0 | 2.5 | 0.37 | -0.9 | 2000 | 10 | 1 | 0.8 | 0.6 | 0.001 | 15 | 9 | 0.31 | 0.56 |
| Oaks | 2 | 27 | 45 | 2.2 | 2.5 | 0.5 | -0.9 | 2000 | 20 | 0.1 | 0.75 | 0.6 | 0.001 | 15 | 9 | 0.21 | 0.59 |

**Table A28.**
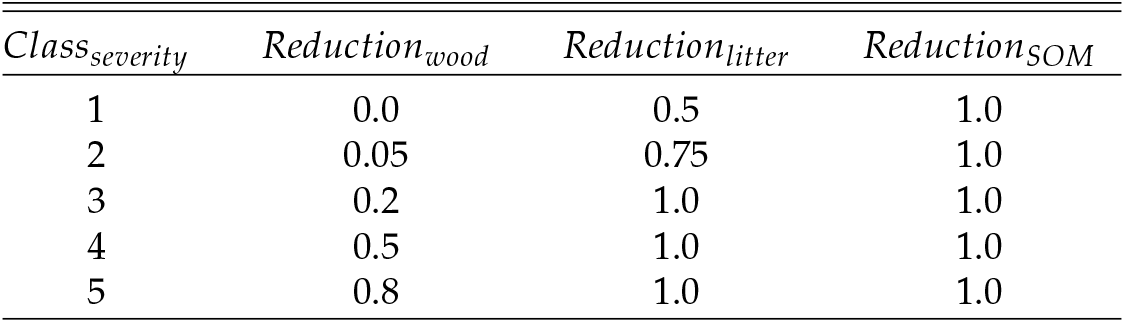
Fire reduction parameters; inactive

**Table A29.** Harvest reduction parameters; inactive

| $Class_{severity}$ | $Reduction_{wood}$ | $Reduction_{litter}$ | $Reduction_{SOM}$ | $Removal_{leaf}$ | $Removal_{wood}$ |
| --- | --- | --- | --- | --- | --- |
| HandThinning | 0.05 | 1.0 | 1.0 | 1.0 | 1.0 |
| MechThinning | 0.05 | 1.0 | 1.0 | 0.85 | 1.0 |

**Table A30.** Species parameters

| Species | Longevity | Maturity | $T_{shade}$ | $T_{fire}$ | $D_{eff}$ | $D_{max}$ | $p_{veg}$ | $S_{min}$ | $S_{max}$ | $R_{fire}$ |
| --- | --- | --- | --- | --- | --- | --- | --- | --- | --- | --- |
| QUIN | 150 | 10 | 4 | 5 | 50 | 3000 | 0.75 | 5 | 40 | resprout |
| QULA | 150 | 20 | 4 | 3 | 50 | 3000 | 0.75 | 5 | 40 | resprout |
| PIPA | 400 | 20 | 1 | 5 | 20 | 200 | 0.0 | 0 | 5 | none |

